# Stimulatrix: An open-source automated platform for high-throughput functional characterization of engineered contractile tissues

**DOI:** 10.64898/2026.09.24.750079

**Authors:** Michael Winkelbauer, Julien Bast, Elena Sophie Casalino, Maria Bulatova, Melanie Generali, Shafeeq Mohammed, Gommar D’Hulst, Francesco Paneni, Katrien De Bock, Thomas Laumonier, Prateek Katiyar, Ori Bar-Nur, Parth Chansoria

## Abstract

Engineered cardiac and skeletal muscle tissues suspended between flexible posts support disease modeling and pharmacology, yet their stimulation, longitudinal imaging and quantitative analysis often remain fragmented and labor-intensive. Here we present Stimulatrix, an open-source platform integrating automated video acquisition within a cell culture incubator, synchronized electrical stimulation and deep learning for longitudinal assessment of contractile tissues in multiwell plates. The platform quantifies tissue compaction, force generation and contraction kinetics with optional cloud processing reducing dependence on local GPU hardware. We demonstrate the workflow in cardiac tissues comprising human induced pluripotent stem cell-derived cardiomyocytes and cardiac fibroblasts, and in primary human skeletal muscle constructs. Force and kinetic measurements were benchmarked against manual annotations. Longitudinal profiling resolved responses to matrix composition and pacing regimens and tracked doxorubicin-associated loss of cardiac contractile force. Stimulatrix provides an accessible workflow for automated functional phenotyping of engineered muscle tissues.

## Main

Human-relevant in vitro models are increasingly being developed as new approach methodologies (NAMs) for mechanistic studies, discovery, and safety and efficacy evaluations.^1^ For contractile tissues like cardiac and skeletal muscle, however, pure physiological relevance depends not only on cellular composition and three-dimensional organization, but also on the capacity to reproduce physiological tissue function.^2^ Genetic perturbations, bioactive compounds and toxicants can alter electrophysiological and contractile phenotypes in muscle tissues, including excitability, contraction and relaxation kinetics, and force generation.^3,4^ Longitudinal functional characterization can therefore complement molecular and structural endpoints by revealing how cellular perturbations translate into contractile phenotypes.^5^

While unconstrained 3D cardiac spheroids offer scalable formats for evaluating tissue formation, multicellular interactions, spontaneous beating and compound responses, they lack a defined axis for force transmission and direct force measurement.^6,7^ This limits their utility for modeling conditions driven by mechanical loading or chronic stress.^8^ In contrast, engineered tissues spanned between flexible posts, develop longitudinal cell alignment and consequently mimic physiological mechanical load and directional force transmission.^9,10^ The repeated displacement of these flexible anchors enables non-destructive estimation of contractile forces and further allows for the assessment of contractile and relaxation kinetics, electrical capture and frequency dependent tissue responses^10,11^. Similar considerations apply to skeletal muscle, where myoblast fusion yields aligned myofibers optimized for directional force transmission. Here, mechanically anchored constructs permit quantitative assessment of twitch, tetanic and fatigue responses under tailored electrical stimulation ^9^. Beyond enabling these measurements, sustained electrical conditioning promotes functional adaptation and tissue maturation, driving key hypertrophic and metabolic shifts, although the resulting phenotype depends strongly on the applied stimulation regimen^12^.

Engineering and functionally characterizing contractile tissues thus involves multiple interconnected steps, from tissue fabrication and electrical conditioning to longitudinal monitoring and quantitative analysis. Existing platforms and computational tools address different components of this workflow: Biowire and Biowire II, for example, combine aligned cardiac tissue formation, electrical conditioning and non-destructive force measurement ^13,14^. Related micropillar arrays such as MyoTACTIC enable the longitudinal characterization of engineered skeletal muscles at higher throughput.^9^ Recently, integrated commercial platforms such as Nautilai,^15^ Mantarray^16^ or Cuore^17^ have enabled repeated or continuous contractility measurements with in-incubator systems using video-based, magnetic and fiber-optic interferometric sensing, respectively. Complementing these experimental platforms, analytical tools including MUSCLEMOTION and BeatProfiler, use image-based analysis to extract contractile, calcium-handling and force-related parameters from microscopy recordings.^18,19^ Despite these advances, currently available solutions either rely on proprietary commercial hardware or address individual components of the workflow, such as stimulation, force sensing or image analysis. An openly accessible system that combines programmable stimulation, in-incubator longitudinal imaging and quantitative analysis within a single hardware–software framework remains lacking. Openly reproducible workflows that integrate tissue fabrication, programmable stimulation, longitudinal optical monitoring and quantitative functional analysis therefore remain scarce. Longitudinal measurements are particularly valuable for modelling progressive disease phenotypes and chronic perturbations, and for assessing interventions whose functional effects emerge over time. Repeated transfer between incubators and microscopes, however, increases handling and exposes tissues to variable acquisition conditions, including changes in temperature and extracellular pH that can alter contractile function ^20^. The primary challenge of engineering is therefore not the absence of individual technologies, but their integration into a robust workflow for repeated stimulation, acquisition and analysis over extended culture periods. Such integration typically requires expertise spanning microfabrication, electronics, automation and computational analysis, which can limit their adoption in laboratories primarily focused on biology, pharmacology or toxicology. A modular approach based on readily available components, openly accessible hardware designs and analysis software, and scalable computational resources could lower this technical barrier while facilitating reproducibility and adaptation across laboratories.

Here, we introduce Stimulatrix, an open-source and modular platform that integrates engineered cardiac and skeletal muscle tissue generation with programmable electrical stimulation, automated in-incubator imaging and quantitative functional analysis. The system is designed around readily available components and openly accessible hardware and software, allowing the complete workflow to be reproduced and adapted without specialized proprietary infrastructure. We demonstrate longitudinal characterization of tissue development and contractile function under spontaneous and electrically controlled conditions. By integrating fabrication, stimulation, acquisition and analysis within a single accessible workflow, Stimulatrix provides a reproducible framework for advanced functional studies of engineered muscle tissues.

## Results

### Design of the Stimulatrix platform for engineered cardiac tissue culture and characterization

To enable rapid acquisition of tissue compaction and contractions over time, we mounted an off-the-shelf digital microscope system (2592×1944 resolution; up to 200x optical zoom) onto an X-Y gantry stage with a 350 mm linear travel along the X axis and 150 mm linear travel along the Y axis (See assembled system in **Figure 2A** and its assembly steps in **Figure S1** and assembled system within the incubator and its associated components in **Figure S2 and Figure S3**). The resulting compact setup could house up to 3 multiwell plates while enabling incorporation in conventional cell culture incubators. Importantly, dynamic culture approaches, including orbital agitation and perfusion, are frequently used in 3D-tissue and organoid culture to enhance mass transport and nutrient exchange.^21^ We therefore designed the plate holder as a mechanically independent module, mounted on an eccentric, motorized stage to provide orbital agitation without interfering with the imaging gantry. For this work, we used 12 well plates for tissue fabrication and culture. The plate lids were modified to accommodate graphite electrodes which were positioned and electrically connected along each plate column using custom 3D-printed housings filled with silver conductive epoxy (**Figure 2B, Figure S4**). One column was left unconnected and served as an unstimulated control. These flexible micropillars were fabricated using reusable molds produced by digital light processing (DLP) printing with a biocompatible resin (**Figure 2C**). Polydimethylsiloxane (PDMS) was cast into the assembled molds under vacuum to ensure complete filling of the pillar features and minimize trapped air. The mold geometry was designed to be readily transferable to alternative manufacturing approaches, including fused-deposition modelling, although DLP-printed molds were used throughout this study. Notably, all the other components of the Stimulatrix platform were 3D printed (see the data availability statement for all CAD files) using Polyethylene terephthalate glycol (PETG) for easy replicability. To further improve replicability and usability, we implemented off-the-shelf electrical components for controlling the motors and applying the stimulation regimen (complete electrical circuitry demonstrated in **Figure S2**; see data availability statement for the code).

**Figure 1.**
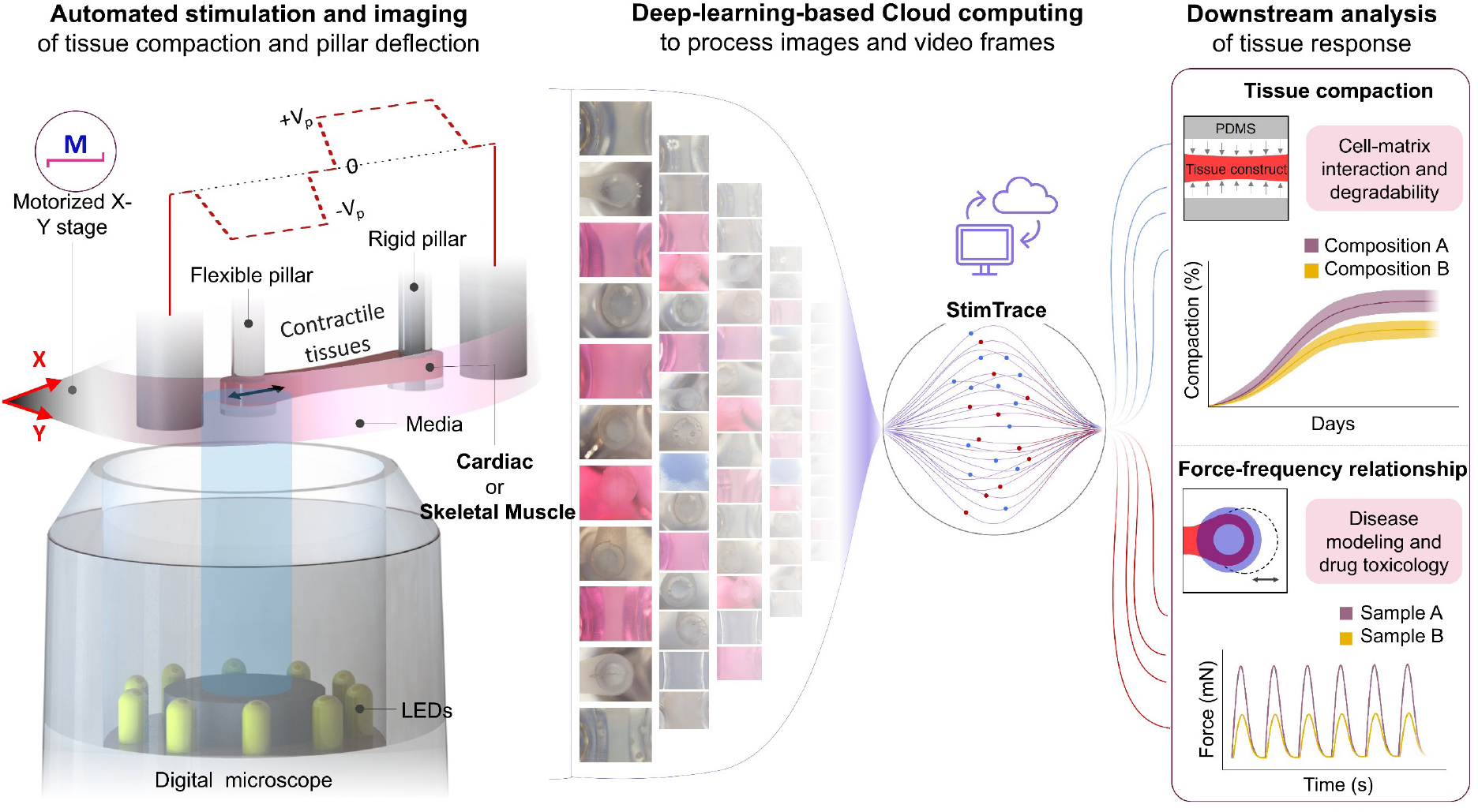
Workflow of the Stimulatrix system and corresponding StimTrace pipeline. Engineered human contractile tissues (cardiac or skeletal muscle) are cast and cultured while being anchored onto two elastomeric pillars (one rigid and another flexible). A digital microscope camera mounted on a motorized stage captures tissue compaction and pillar deflections – with or without electrical stimulation – over time. The image frames are fed through a deep learning pipeline which automatically plots tissue compaction and contraction metrics for downstream analysis of different training regimen or material and drug responses.

**Figure 2.**
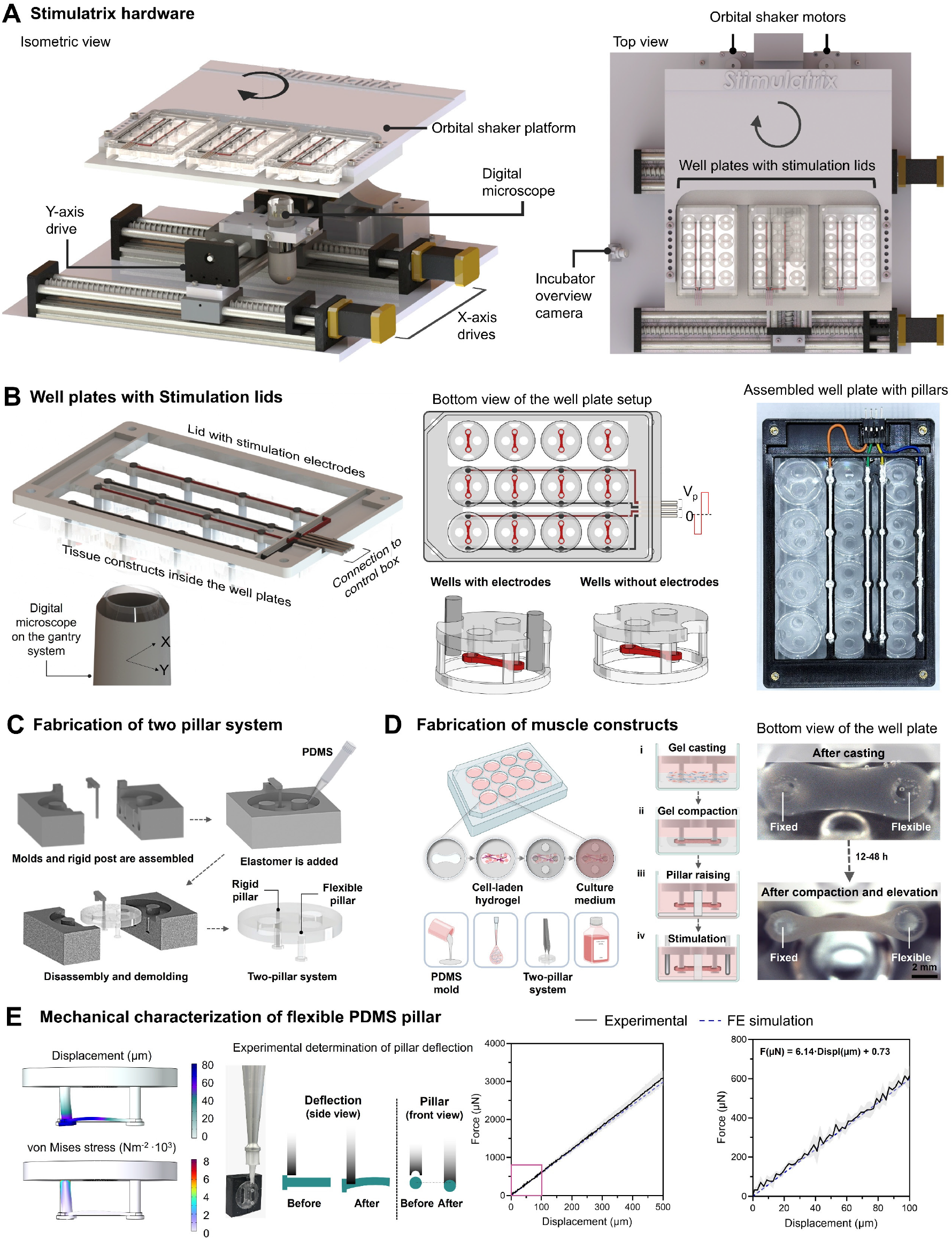
Stimulatrix system and its physical components. **A.** Isometric and top views of the system hardware demonstrating X-Y gantry and digital microscope used to image tissue compaction and contraction from below. Tissue culture plates are positioned on an orbital shaker platform above the camera, which accommodates up to three well plates. **B.** Stimulation lids with corresponding electrical connection for graphite electrodes. Two rows are stimulated in quadruplicates, while the third row without the electrodes can be used for unstimulated controls. **C.** Fabrication of pillar molds involves casting PDMS (1:15 crosslinker ratio) between two negative molds with a 3D printed rigid pillar pins inserted in the slot for casting one rigid pillar. **D.** Bioresins (cells+matrix components; details in the results and methods) are cast into well plates with the casting molds and the pillar constructs are inserted into the wells. After compaction over 3 days, the constructs are raised from the pillar platform. **E.** The pillar deflection and corresponding forces are determined through computational models, and subsequently validated through bending force measurements, yielding a close correlation between the analytical and actual forces and enabling reliable force estimation from captured pillar displacement.

The tissue constructs were fabricated by gently mixing human cells – iPSC-derived cardiomyocytes and cardiac fibroblasts or myogenic progenitor cells – with fibrinogen supplemented with Matrigel^®^ (details in methods) and casting in customized plates containing dumbbell-shaped PDMS wells (**Figure S5**). The cell-matrix mixture underwent compaction over time (**Figure 2D**), and the constructs were subsequently raised using 3D printed inserts, freeing up the flexible posts to undergo bending. Importantly, we also computationally modeled and experimentally validated the mechanics of the pillars to determine a relationship between obtained auxotonic contraction amplitude and the forces generated by the muscle constructs to deflect the pillar (**Figure 2E**).

### Deep-learning-based analysis of tissue compaction and contractility

The Stimulatrix system was controlled using a custom software which enabled image and video acquisition of selected wells at the desired intervals, and electrical stimulation at selected regimen (see **Video S1**). At different phases of the culture period, the gantry moved underneath the desired well and captured: **1.** Snapshots at the center of the dumbbell shaped tissue casting mold to characterize the tissue compaction over time, and **2.** Videos of pillar deflections to characterize spontaneous tissue contractions (for cardiac constructs) in response to the electrical stimulation (for both cardiac and skeletal muscle constructs). The captured images and videos were then fed into a deep learning pipeline (“Stimtrace”) based on the widely utilized U-net architecture (pipeline illustrated in **Figure 3A**). To circumvent the need for dedicated local GPU hardware, model development and analysis were initially implemented using Google Colab with Google Drive for data storage, providing browser-based access to the complete pipeline. The analysis workflow was additionally configured for local execution, allowing users to select computational resources according to availability. During model development, geometric and photometric tranforms applied to the acquired image data using the *Albumentations* library augmented the training dataset and improved tissue and pillar segmentation losses (**Figure 3B**). For the pillar segmentations, addition of ellipse fit algorithm further reduce the total validation loss, yielding consistent pillar prediction outputs with minimal jitter when the pillar was static. To benchmark StimTrace against a conventional computer-vison approach, we compared the deep-learning segmentation pipeline with Lucas-Kanade optical-flow based point tracking (see methods). Under high-contrast imaging conditions, both approaches reliably captured pillar displacement (see comparison of the different methods in **Figure S6,** and the corresponding segmentation workflow in **Video S2** and point tracking in **Video S3)**. However, the performance of optical-flow tracking deteriorated under imaging conditions more representative of longitudinal culture, including changes in medium color, non-uniform illumination, optical reflections, jitter and the presence of cellular debris. These perturbations increased noise and feature loss, resulting in tracking drifts and disproportionately affecting derivative-based parameters such as contraction and relaxation rates. In contrast, StimTrace maintained stable pillar segmentation across these conditions, yielding more consistent force-frequency traces and contractility metrics (see force-frequency traces in **Figure S6,** and overlayed recording in **Video S5**). These results demonstrate that segmentation-based pillar tracking improves robustness to imaging variability typically encountered during longitudinal culture.

**Figure 3.**
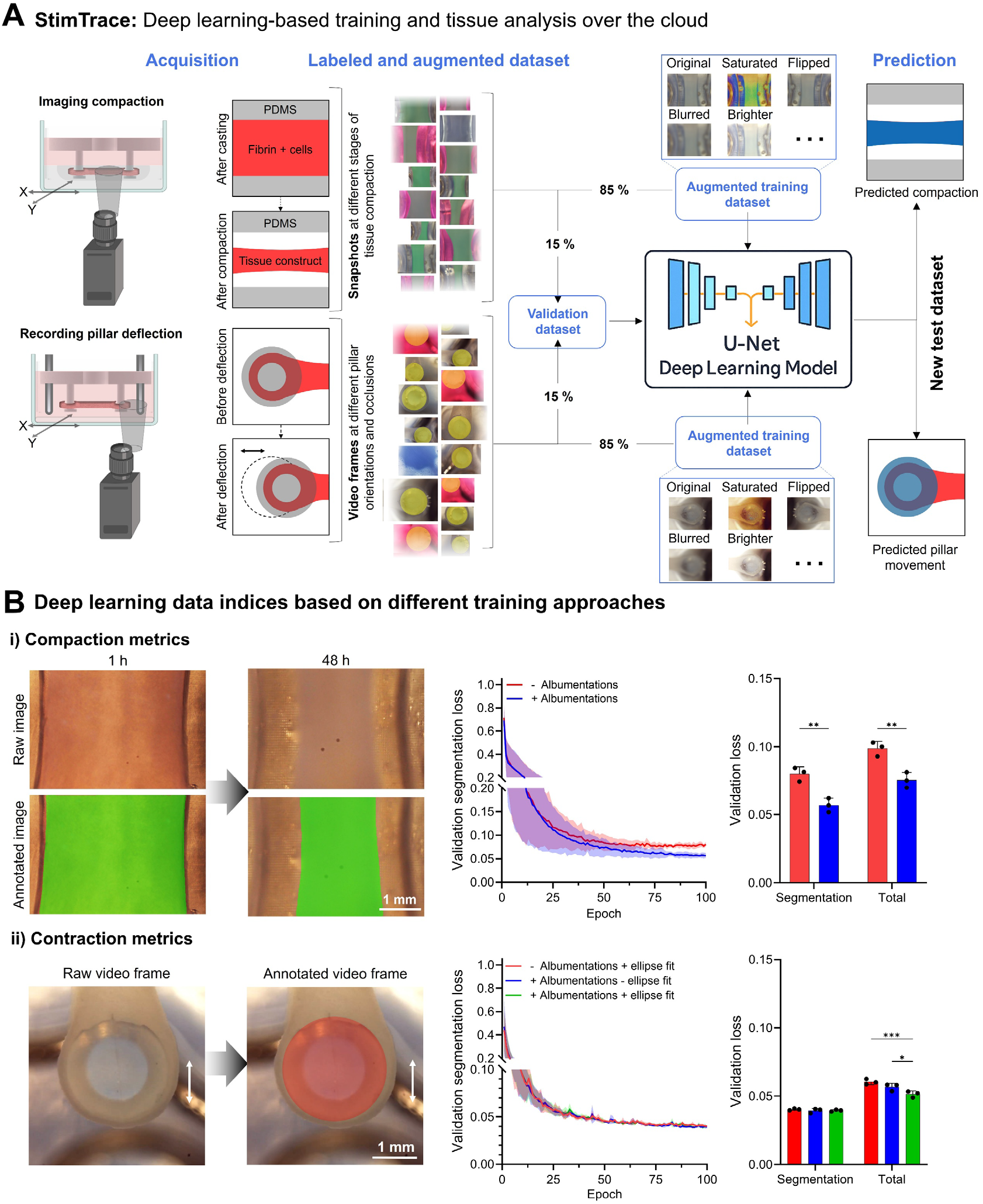
StimTrace pipeline and its optimization for robust analysis of tissue compaction and contraction dynamics. **A.** The acquired images (i.e., recorded tissue compaction over time) or videos (i.e., recorded pillar deflection due to tissue contractions) fed into the StimTrace pipeline. A pre-selected dataset – with manual traces of the tissues (in compaction snapshots) or pillars (on contraction videos) – is augmented via Albumentations library and used for training the deep learning (U-net) pipeline to predict tissue compaction and pillar deflection. **B.** As the training progressess, the segmentation and total loss for the validation set for both compaction (subpanel i) and pillar deflection (subpanel ii) is lower when Albumentations library is used to augment the training dataset. Additionally, for the pillar deflection (subpanel ii), using an ellipse fit post-processing in the dataset further reduces the total validation loss. Bar plots show the total validation loss at the end of 100 training epochs.

### Automated assessment of tissue compaction and contractility enables cell and matrix optimization

The Stimulatrix system with the subsequent StimTrace pipeline enabled longitudinal assessment of the matrix compaction tissue mechanics under different cell and matrix compositions. For the cardiac constructs, we investigated different concentrations of fibrinogen (3 and 5 mg/mL; referred to as fibrin after crosslinking) and cell compositions (only iPSC-derived cardiomyocytes, and further addition of cardiac fibroblasts at different ratios, while keeping the total cell concentration constant (10^7^ cells/mL) (**Figure 4A**). For the skeletal muscle constructs, we only used one cell type (primary human myogenic progenitor cells; Pax7^+^; 8·10^6^ cells/mL) but changed the matrix composition to have different concentration of fibrinogen (3, 5 mg/mL) including Matrigel^®^ (30% v/v). For both tissue types, % compaction (based on the width of the segmented images after the StimTrace pipeline was applied) was tracked for up to 48 after casting. For the cardiac constructs, matrix compaction was greater at lower fibrin concentration (3 mg/mL) and when cardiac fibroblasts comprised 25% of the total cell population,^22^ consistent with previously established formulations^22^. Lower fibrin concentrations facilitate greater matrix densification (likely due to less dense and mechanically resistant fibrin networks). The addition of cardiac fibroblasts further increased compaction, consistent with their capacity to adhere to, remodel and contract fibrillar extracellular matrices through integrin-mediated actomyosin traction.^23^ For the skeletal muscle constructs, lowering the fibrin concentration demonstrated similar compaction characteristics as the cardiac constructs. The addition of Matrigel® at 30% v/v further increased matrix compaction,^24^ potentially reflecting both reduced mechanical resistance of the composite matrix and enhanced cell-matrix interactions that facilitate myoblast-mediated matrix remodeling. Overall, matrix compaction rates generally plateaued over 48 h, which was the time-period selected for elevating the constructs prior to further maturation and contraction response evaluation.

**Figure 4.**
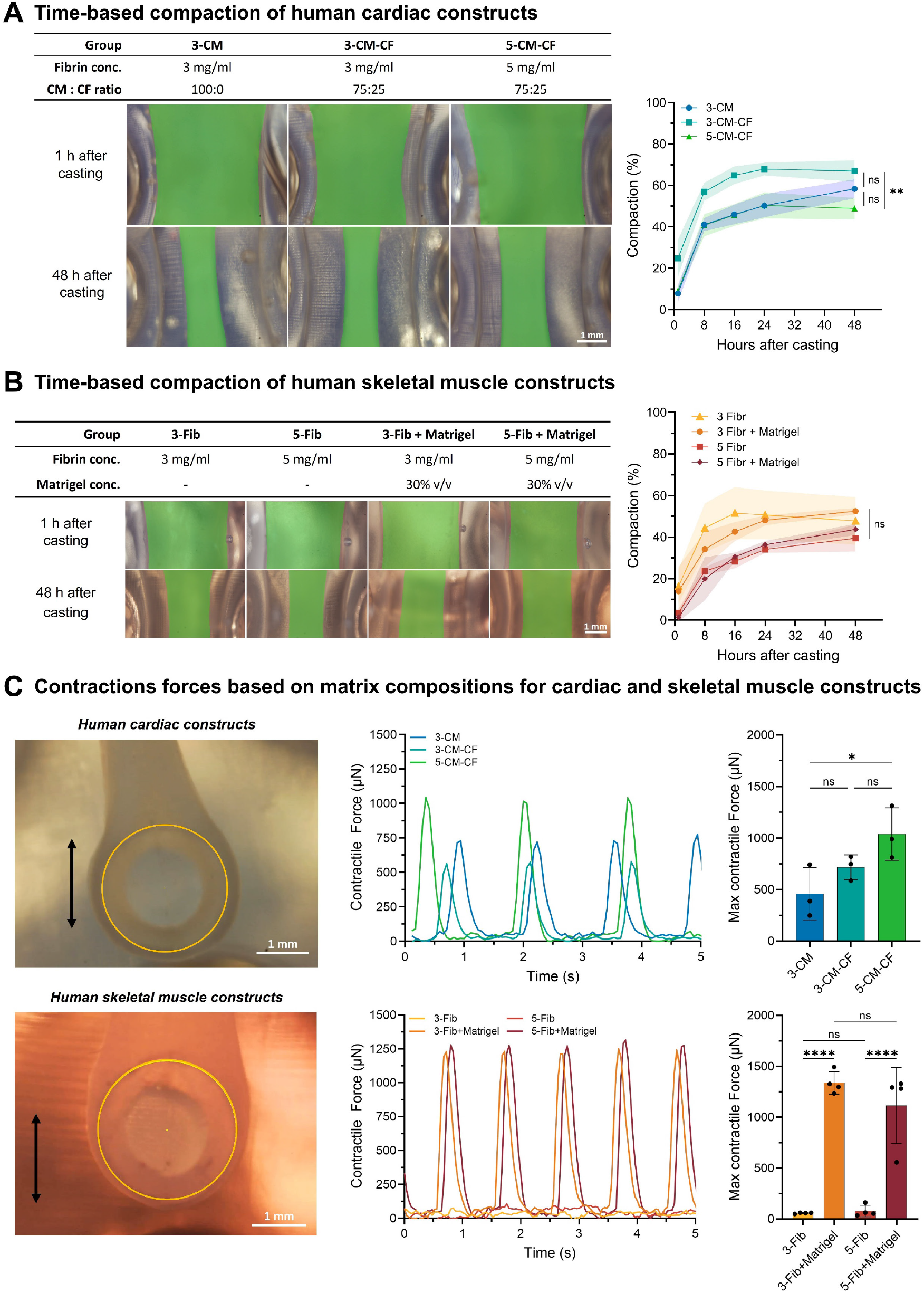
Stimulatrix platform enables autonomous tracking of various cell and matrix compositions of human cardiac and skeletal muscle tissues. **A.** Longitudinal tracking of the compaction of human cardiac constructs featuring a combination of iPSC-derived cardiomyocytes (CMs) and cardiac fibroblasts (CFs) at different mixing ratio demonstrates that higher fibrin concentration leads to lower matrix compaction, while higher number of cardiac fibroblasts leads to higher compaction in the matrices. Of note, the total cell concentration was kept the same (10^6^ cells/mL). Scale bar: 1 mm. **B.** For human skeletal muscle constructs, although lower fibrin concentration demonstrated higher compaction and the addition of Matrigel did not significantly affect the compaction metrics. Scale bar: 1 mm. **C.** Longitudinal tracking of the contraction metrics enables material optimization for both cardiac and skeletal muscle constructs. Here, a 5 mg/mL fibrin only matrix together with a combination of CMs and CFs results in the highest contraction forces for cardiac constructs, while for skeletal muscle constructs the forces are highest for 3 mg/mL fibrin with 30% v/v Matrigel with encapsulated myogenic progenitor cells. Data represented as mean ± SD (n >= 3), statistical significance was determined by one-way ANOVA with Tukey’s post-hoc analysis and is denoted as follows: **** represents p < 0.0001, * represents p < 0.05, ns = not significant.

The contraction responses were evaluated for both cardiac and skeletal muscle constructs after 2 weeks of maturation. For the cardiac constructs, no electrical stimulation was applied when mapping the spontaneous contractile responses. In contrast, skeletal muscle constructs did not demonstrate any spontaneous contractility. Here, following 5 days of maturation of static culture (i.e.: without electrical stimulation), the constructs were subjected to biphasic 1-Hz electrical pacing regimen (for 6 min, once every hour) using the Stimulatrix stimulation software (see data and source code availability statement for details) before functional assessment. Notably, charge-balanced biphasic stimulation was used to minimize Faradaic reactions at the electrode-medium interface, thereby reducing electrochemical degradation during prolonged stimulation. The StimTrace pipeline enabled robust segmentation of the elastomeric pillars and tracing of the centroid displacement (**Figure 4A**), which was converted to contraction forces using the force-displacement relationship previously derived (**Figure 2E**). For the cardiac constructs, contractile forces were significantly greater in tissues containing both cardiomyocytes and cardiac fibroblasts than in cardiomyocytes-only constructs. This may reflect fibroblast-mediated matrix remodeling and compaction, which can improve tissue organization and mechanical coupling between cardiomyocytes. Previous fibrin-based engineered cardiac tissue studies similarly reported enhanced tissue compaction and contractile performance following the addition of cardiac fibroblasts.^25^ By contrast, fibrin concentration did not significantly affect contractile force within the tested range. Nevertheless, constructs containing 5 mg/mL fibrin were mechanically more robust and less prone to rupture than those containing 3 mg/mL fibrin, justifying the higher concentration for subsequent experiments. In skeletal muscle constructs, addition of Matrigel® to the fibrin matrix substantially increased contractile force, whereas fibrin concentration itself did not significantly affect force generation, consistent with the cardiac constructs. We therefore selected 5 mg/mL fibrin for subsequent experiments because it provided greater mechanical robustness without compromising contractile performance.

The fully automated high-throughput screening using the Stimulatrix system enabled rapid screening and optimization of the material combination to boost the contraction forces. Here, a combination of cardiomyocytes and cardiac fibroblasts (75:25 ratio; total concentration 10^7^ cells/mL) and matrix composition of fibrin at 5 mg/mL was deemed optimal for cardiac constructs, while human skeletal muscle constructs required the addition of Matrigel^®^ at 30% v/v to obtain high contraction forces. These compositions were used for all subsequent studies.

### Stimulatrix enables controlled electrical pacing and maturation of engineered cardiac tissues

When designing the Stimulatrix components, we decoupled the tissue-anchoring pillar inserts from the well plate to enable free rotation of the micropillars (on which the tissues were anchored) inside the wells. This enabled changing the orientation of the engineered tissues relative to the electrodes, resulting in either an orthogonal or collinear electrical field orientation through the tissues (**Figure 5A**). Here, computational modelling showed that the collinear configuration generated an electric field predominantly aligned with the tissue long axis, with an approximately linear potential gradient along this direction. In contrast, the orthogonal configuration generated an approximately uniform electric field transverse to the tissue long axis, while the electric potential remained nearly constant along the length of the tissues. We further investigated the impact of these electrode configurations on tissue contractions. Cardiac constructs were used to assess electrical capture because their spontaneous contractile activity provided a baseline against which stimulation-induced pacing could be distinguished. Prior to testing, the tissues underwent progressive electrical conditioning, with the stimulation frequency increased from 1.5 Hz to 5 Hz over the culture period of 14 days (**Figure 5C**). Following this conditioning regimen, the tissues synchronized to the applied pacing frequency and maintained capture at frequencies of up to 5 Hz in both electrode configurations (**Figure 5B**). Tissues maintained 1:1 mechanical capture up to 5 Hz, without obvious alternans or arrhythmic contractions. Furthermore, increasing stimulation frequencies progressively reduced contractile force, consistent with the negative force-frequency relationship commonly observed in immature hiPSC-derived cardiac tissues. This response has been attributed to incomplete maturation of intracellular Ca^2+^ handling, particularly limited sarcoplasmic-reticulum Ca^2+^ cycling at higher pacing frequencies.^26^ When comparing the two electrode orientations no difference was observed in terms of the contraction forces at continuous or ramped frequency stimulation (**Figure S7**). For all subsequent experiments, collinear electrode orientation was used because it aligned the applied electric field with the tissue long axis and at the same time facilitated routine medium changes more easily. Interestingly, cardiac constructs undergoing electrical conditioning displayed significantly longer sarcomere lengths than unstimulated controls, measuring 2.06 ± 0.15 µm for Collinear and 2.04 ± 0.18 µm for Orthogonal configurations versus 1.93 ± 0.17 µm for Control (**Figure 5D**). Similar trends were observed for the gap junction protein Connexin 43 (Cx43), where electrical pacing led to both a higher number of Cx43 puncta (Figure 5D) and a significant increase in puncta spot size (**Figure 5D, Figure S8**).

**Figure 5.**
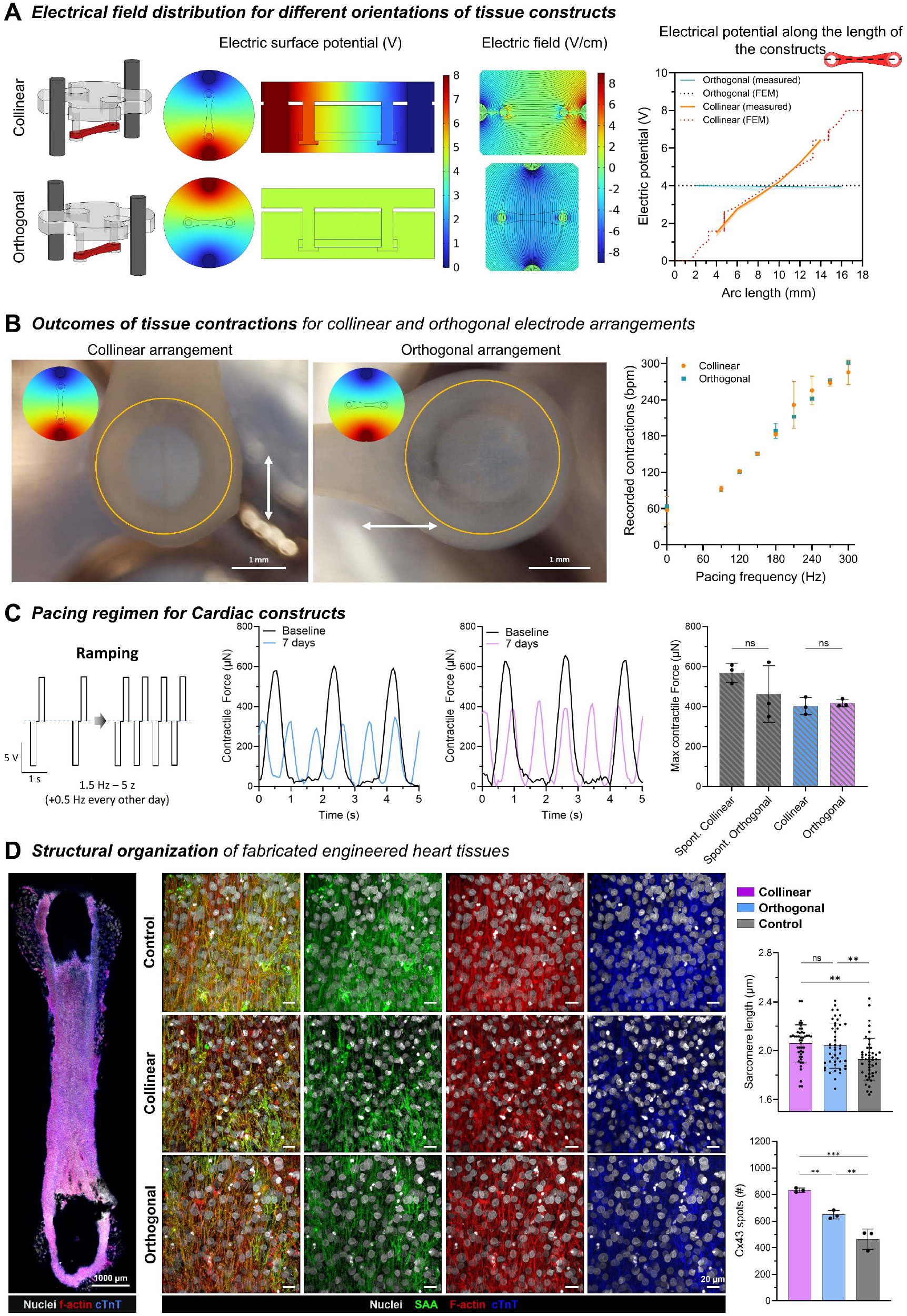
Stimulatrix system enables facile changes in electrical field orientation and subsequent contraction dynamics tracking. **A.** By simply rotating the pillar arrays within the well plates, the electrical field orientations can be changed. Computational models demonstrate electrical field lines either collinear with muscle axis, or perpendicular to the muscle axis. Experimental validation of the electric potential (right) also demonstrates close correlation with the computational model, with an approximately linearly increasing electric potential observed in the collinear electrode arrangement, while a constant field is observed in the orthogonal arrangement. **B.** Video acquisition by the Stimulatrix system and subsequent pillar segmentation using the StimTrace pipeline enables tissue contraction assessment when oriented parallel or orthogonal to the electric field. The tissues demonstrated synchronicity with applied electrical stimulus under both field orientations. **C.** Long-term electrical pacing regimen in cardiac tissues. Schematic of the progressive frequency-ramping protocol (1.5 to 3 Hz over 7 days) with pre-conditioned and day 7 force-traces, respectively. Bar chart shows maximum developed contractile force across spontaneous and paced conditions in collinear vs. orthogonal configuration. **D.** (Left) Structural characterization and sarcomeric maturation of cardiac tissue following electrical conditioning. Stitched confocal overview of cardiac constucts morphology (scale bar 1000 µm), (Middle) High magnification immunostaining for nuclei (grey), SAA (green), F-actin (red), and cardiac troponin T (cTnT, blue) (scale bars: 20 µm). (Right) Quantitative analysis of sarcomere length and Connexin 43 (Cx43) spot density across conditions. Data represented as mean ± SD (SAA length: n = 45, Cx43(#): n=3), statistical significance was determined by one-way ANOVA with Tukey’s post-hoc analysis and is denoted as follows: *** represents p < 0.001, ** represents p < 0.01, ns = not significant.

Quantitative orientation analysis of F-actin filaments revealed distinct structural responses across pacing regimens (**Fig. S9**). In engineered heart tissues, unpaced control samples exhibited the highest degree of longitudinal myofibrillar alignment (83 ± 1.8%), which was significantly greater than in orthogonal (69 ± 6.1) and collinear conditions (54 ± 3.7%). Orthogonal pacing also preserved significantly higher alignment than collinear pacing Conversely, engineered skeletal muscle tissues demonstrated consistently high structural orientation across all conditions, showing no statistically significant differences in F-actin alignment among continuous, pulsed, and control groups (**Fig. S9B**).

### Programmable electrical conditioning enhances functional and structural maturation of engineered skeletal muscl

To enable customizable electrical conditioning of engineered skeletal muscle – where twitch, summation, and tetanic responses require distinct pacing profiles (**Figure 6A**) – tissues were subjected to distinct training protocols following 5 days of static culture. Constructs were exposed to either continuous 1 Hz stimulation (6 min/h) to simulate endurance-like training or 10 Hz pulse trains (9 s rest intervals, 6 min/h) to model high-intensity interval training. Longitudinal force monitoring revealed a time-dependent increase in contractile force across both conditioning groups (**Figure 6A**), with 10 Hz stimulation reliably eliciting high-amplitude tetanic contractions. After 7 days of conditioning, functional adaptation was evaluated relative to unpaced static controls. Continuously trained tissues and controls were acutely evaluated at 1.5 Hz, while high-intensity trained tissues and controls were tested using 10 Hz pulse trains. Both electrical conditioning regimens significantly improved functional performance, yielding higher contractile forces than unstimulated controls. To determine whether these functional gains correlated with structural remodeling, tissues were immunostained (**Figure 6B**) for myosin heavy chain (MyHC) and sarcomeric-α-actinin (SAA). Consistent with force measurements, electrical conditioning significantly enhanced sarcomeric maturation, yielding average sarcomere lengths of 2.43 ± 0.15 µm for continuous, 2.27 ± 0.13 µm for pulsed, and 2.04 ± 0.21 µm for control tissues. Although not statistically significant, similar positive trends were observed in myotube diameter and fusion index, with continuously stimulated constructs displaying the thickest myotubes and highest degree of fusion.

**Figure 6.**
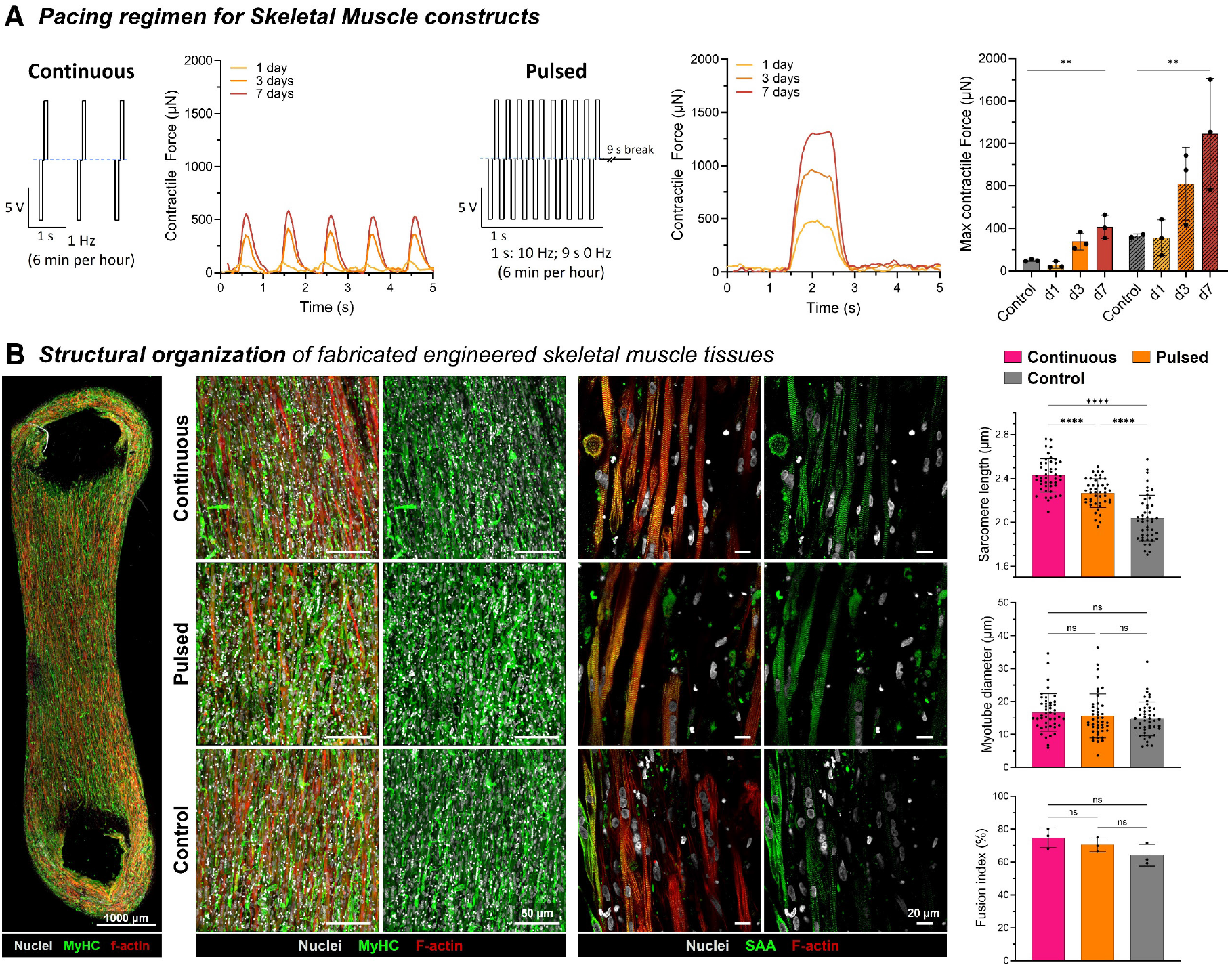
Controlled training and structural organization of skeletal muscle constructs. **A.** Pacing regimens for skeletal muscle tissue constructs. Following 5 days of static culture, tissues were subjected to either continuous 1 Hz stimulation (6 min/h, solid bars) to simulate endurance-like training or 10 Hz pulse trains (with 9 s rest intervals, 6 min/h, hatched bars) to model high-intensity training. **B**. (Left) Stitched confocal overview of engineered skeletal muscle tissue morphology (scale bar: 1000 µm; stained for nuclei, grey; myosin heavy chain (MyHC), green; F-actin, red). (Middle) High-magnification confocal micrographs showing immunostaining for nuclei (grey), MyHC (green), and F-actin (red; scale bar: 50 µm), alongside nuclei (grey), sarcomeric α-actinin (SAA, green), and F-actin (red; scale bar: 20 µm) across conditions. (Right) Quantitative analysis of structural parameters: continuous electrical pacing significantly enhanced sarcomeric maturation (sarcomere length) compared to high-intensity pacing and unpaced controls, whereas no significant differences were observed for myotube diameter or fusion index. Data represented as mean ± SD (SAA length: n = 45, Myotube diameter n=45, Fusion index: n = 3). Statistical significance was determined by one-way ANOVA with Tukey’s post-hoc analysis and is denoted as follows: **p<0.01, ****p < 0.0001, ns = not significant).

### Automated functional profiling captures drug-specific cardiac responses and chronic cardiotoxicity

Having established that the Stimulatrix platform and the StimTrace deep-learning pipeline enable rapid biomaterial optimization and precise force tracking across both human skeletal and cardiac muscle constructs, we next sought to demonstrate the platform’s utility in automated safety pharmacology. Although skeletal muscle tissues allow robust characterization of twitch and tetanic force kinetics under defined stimulation regimens, the spontaneous electromechanical activity of human engineered cardiac tissues permits unpaced, non-destructive monitoring of both beating rate (chronotropy), relaxation kinetics (lusitropy) and force generation (inotropy) over extended culture periods. Because cardiovascular toxicity represents one of the primary drivers of pharmaceutical compound failure, we leveraged the automated incubation, multi-well imaging, and cloud-based segmentation capabilities of Stimulatrix to perform multi-parametric pharmacological and toxicological screening on optimized cardiac constructs. Here, we evaluated a panel of five reference therapeutics selected to probe orthogonal molecular pathways regulating cardiac excitation–contraction coupling, sarcolemmal ion fluxes, myofilament calcium handling, repolarization dynamics, and chronic cardiotoxic exposure. Acute administration of the *β*-adrenergic agonist isoproterenol (10 - 1000 nM)^27^ elicited a concentration-dependent positive chronotropic response and accelerated contraction/relaxation kinetics (referenced to baseline levels; **Figure S10**), characterized by reduced time-to-peak (TTP), contraction duration (CTD_50_), and relaxation time (Relaxation_50_), reflecting expected positive lusitropy while maintaining peak force generation (**Figure 7A**, **Figure S10**). The limited inotropic response to isoproterenol and negative force–frequency relationship likely reflect the immature electrophysiological and calcium-handling phenotype characteristic of hiPSC-derived cardiomyocytes, rather than mature adult-like myocardial physiology.^26,28^ Similarly, the calcium sensitizer levosimendan (30 - 3000 nM)^27^ increased beating rate and shortened contraction kinetics (CTD_50_, CTD_90_ and Relaxation_50_, Relaxation_90_), accompanied by a dose-dependent reduction in mean contractile force at the highest concentration (3000 nM; **Figure 7B**, **Figure S10**). Conversely, blocking L-type calcium channels with nifedipine (30–300 nM)^27^ induced severe, dose-dependent negative inotropy, reducing contractile force by > 80% at 300 nM alongside marked decreases in maximum contraction and relaxation slopes and a reduction in beating frequency (**Figure 7C**, **Figure S10**). Assessment of the antiarrhythmic drug sotalol (3–30 µM)^29^ demonstrated maintenance of peak force amplitude combined with concentration-dependent prolongation of contraction duration (CTD_50_, CTD_90_) and reduced beating frequency (**Figure 7D**, **Figure S10**). These mechanical responses are compatible with the established electrophysiological actions of sotalol, although action-potential and calcium-transient measurements were not performed and therefore the underlying mechanism cannot be resolved from the contractile readouts alone.^30^ Finally, to demonstrate the utility of the system for chronic cardiotoxicity modeling, constructs were exposed to doxorubicin. Dose-response screening across 48 h in 3D cardiac spheroids established an IC50 (half-maximal inhibitory concentration required to reduce the cell viability by 50% relative to untreated controls) of 4.5 µM (**Figure S11**), identifying 1 µM as a mildly cytotoxic concentration that maintains short-term metabolic viability. Upon extended treatment with 1 µM doxorubicin over 96 h, longitudinal monitoring revealed a progressive, time-dependent decline in maximum contractile force, becoming statistically significant by 72–96 h relative to vehicle controls (**Figure 7E**). Together, these distinct functional and kinetic profiles confirm that the platform faithfully recapitulates physiological inotropic, chronotropic, lusitropic, and chronic toxicological responses.

**Figure 7.**
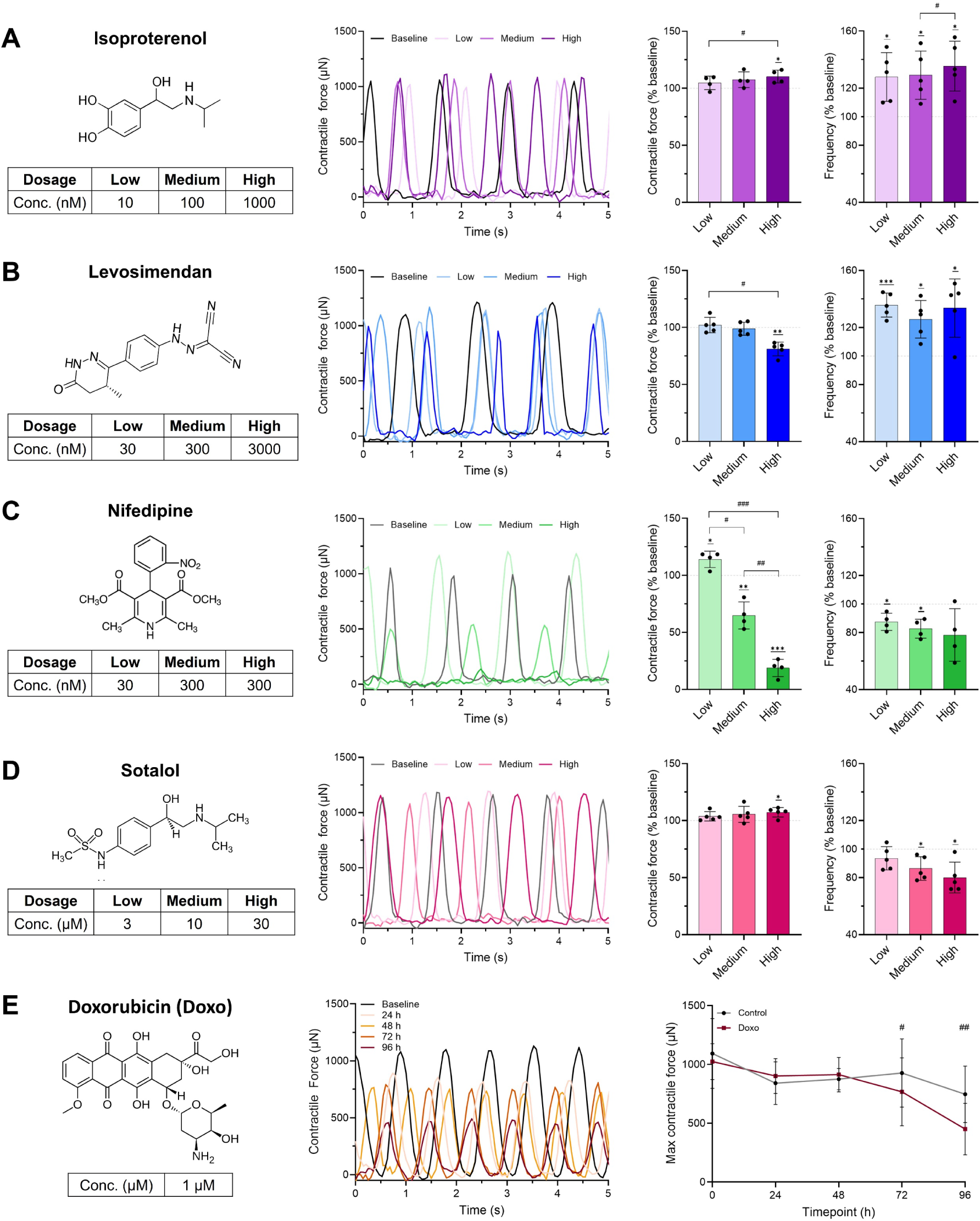
Automated in vitro toxicology on engineered cardiac tissues exposed to selected off-the-shelf therapeutics at different concentrations (in the media). **A. Isoproterenol**, a β-adrenergic agonist expectedly increased the contraction frequency, but did only affect the peak contractile force in the highest concentration, likely due to the immature or early-stage phenotype of human engineered cardiac constructs which typically show a weak or negligible inotropic response. **B. Levosimendan**, a calcium sensitizer and PDE3 inhibitor, increased beating frequency while exhibiting a drop in force generation at high concentrations, likely due to off-target PDE3 inhibition or saturation of intracellular calcium handling**. C. Nifedipine**, an L-type Ca^2+^ channel blocker, induced a severe, dose-dependent decrease in contractile force accompanied by a reduction in beating rate, reflecting the primary dependence of engineered cardiac tissues on extracellular calcium influx. **D. Sotalol**, a Class III antiarrhythmic (K^+^ channel blocker), reduced beating frequency and significantly prolonged contraction duration while maintaining peak contractile force, consistent with expected repolarization delay. **E. Doxorubicin** (Doxo) a cardiotoxic chemotherapeutic agent, caused a progressive, time-dependent reduction in contractile force over 96 h of exposure (1 µM), confirming the platform’s capacity to detect chronic, sub-lethal functional cardiotoxicity over extended culture periods. Data represented as mean ± SD (n = 4-5 independent constructs per group). Statistical significance was determined by one-way ANOVA with Tukey’s post-hoc analysis. Significance relative to baseline (100%) is denoted by asterisks (*p < 0.05, ** p < 0.01, *** p < 0.001), while significance between groups is denoted by hash marks (# p < 0.05, ## p < 0.01, ### p < 0.001).

## Discussion

The Stimulatrix platform unites mechanized video acquisition, customizable electrostimulation, and deep learning-based automated segmentation into an open-source, in-incubator workflow for cultivation and functional characterization of engineered cardiac and skeletal muscle tissues. Quantifying microtissue contractility via micropillar deflection has conventionally relied on optical-flow tracking methodologies demonstrated by Lucas–Kanade^31^ or Horn and Schunck.^32^ However, optical flow performance degrades significantly during extended tissue culture due to non-uniform illumination, media color shifts, optical reflections, frame jitter, and cellular debris shedding. The deep learning-based StimTrace pipeline, leveraging an augmented U-Net framework, overcomes these imaging artifacts by learning structural boundaries rather than relying on pixel-intensity gradients (see overlayed recording in **Video S5**, and force-frequency traces in **Figure S6)**. Future studies could investigate hybridization of the deep learning and optical flow tracking for robust assessment of contractile kinetics.^33^ For instance, the deep learning-based segmentations can enable detection of pillars of various shapes and sizes (including different lighting and color conditions) and optical flow tracking can then track pillar movements.

For the first set of cell experiments, the platform enabled rapid screening of cell and matrix compositions for boosting contraction forces. In this study, while Matrigel was omitted from cardiac constructs to maintain a simplified, defined bioresin formulation, adding 30% v/v Matrigel to the fibrin matrix proved essential for achieving high contractile forces in skeletal muscle constructs, likely due to enhanced basement membrane cues promoting myoblast alignment and fusion. Notably, incorporating basement membrane components such as Matrigel, fibronectin or laminin into cardiac bioresins could further boost baseline contractile forces and structural alignment.^34,35^ In the cardiac constructs, addition of fibroblasts significantly enhanced matrix compaction and maximum active force compared to cardiomyocyte-only tissues, aligning with established roles of fibroblasts in integrin-mediated matrix remodeling and mechanical coupling.^25^ Here, incorporating endothelial cells and pericytes could further establish microvascular networks, improving mass transport, paracrine signaling, and physiological force transmission in thicker tissue constructs.

In our experiments, progressive electrical pacing regimens successfully promoted structural maturation across both tissue types, significantly increasing sarcomere length (>2.4 µm in skeletal muscle and >2.0 µm in cardiac constructs) and upregulating gap-junction Connexin 43 spot size and density (**Figure S8**). During extended electrical conditioning, longitudinal shifts in electrode impedance and culture media conductivity can alter effective charge injection over time. Although charge-balanced biphasic pulses reduced net electrode polarization and electrochemical degradation in our setup, integrating real-time, closed-loop impedance and current sensing into future stimulator iterations could dynamically adjust voltage outputs to maintain constant current delivery across multi-week pacing protocols. Because the primary objective of the present study was to establish and validate the foundational platform technology, exhaustive environmental and pacing permutations would be a future scope of investigation. Accordingly, future studies will systematically deploy the platform to evaluate long-term tissue maturation timelines, specialized electrical conditioning protocols (e.g., chronic frequency ramps or high-intensity interval trains),^12,36^ and combined electromechanical conditioning by coupling electrical stimulation with dynamic mechanical loading via the integrated orbital stage, by changing the stiffness of the elastomeric pillars, and introducing remote straining capabilities into the system.^26^

A current limitation of the platform is that validation was performed in a 12-well format, with relatively small numbers of biological replicates. Although the modular design should permit adaptation to 48- or 96-well formats, higher-density formats introduce several technical challenges. First, because miniaturized tissue constructs are expected to generate lower absolute contractile forces, accurate camera-based measurement requires the optimization of the elastomeric pillar geometries or using softer formulations to maintain detectable deflections. Second, the spatial constraints of positioning dense electrode pairs in close proximity complicate conventional lid fabrication, necessitating alternative strategies such as custom microelectrode arrays^16,37^ or optogenetic stimulation^38,39^ for tissue pacing. Third, the serial nature of gantry-based video acquisition becomes increasingly time-consuming across high-density plates; this can be overcome by incorporating wide-field telecentric optics or camera arrays to capture high-resolution pillar deflections across multiple wells simultaneously.

In this work, we established independent workflows for the Stimulatrix and StimTrace software workflows, as that enables each platform to be easily tuned and implemented to new well plate formats and tissue construct geometries. For instance, once the hardware is established, the Stimulatrix platform software could be used for controlled stimulations and video acquisitions in selected wells, regardless of the design of the well plate inserts, pillar arrays and geometry of the muscle constructs. Similarly, once the videos or snapshots of the tissues are acquired – through the Stimulatrix platform or any other setup such as a microscope with gantry-based imaging system - the StimTrace pipeline could be used for contractility and compaction analysis after appropriate augmentation of the training pipeline. Similar inbuilt analytical analysis tools including optical flow-based point tracing are available in the StimTrace workflow and could be used provided sufficient image quality and minimal perturbations persist when performing the video acquisitions. Importantly, labs replicating our work could also deploy a unified pipeline for video acquisition and analysis in the same software to simplify the workflow.

Practical hardware and computational constraints also present specific operational trade-offs that can be addressed in future development. Executing StimTrace on cloud platforms (e.g., Google Colab) eliminates local GPU hardware barriers for biology laboratories, but uploading large longitudinal video datasets to cloud servers introduces data protection, bandwidth, and latency challenges. This can be mitigated by performing model inference locally on consumer-grade hardware, such as standard desktop GPUs or gaming PCs, or by deploying optimized model compression techniques (e.g., TensorRT quantization) directly on local edge compute units.

## Materials and Methods

### Deep Learning training pipeline for pillar segmentation (contraction metrics)

To eliminate local GPU hardware constraints and maximize accessibility, the StimTrace software (written in Python) interfaced with Google Colab (integrated directly with Google Drive for cloud storage) for model training and checkpointing. The dataset for nodel development was partitioned into disjoint sets with 85% samples in the training set and 15% samples in the validation set. Input images and manually labeled ground-truth masks were standardized to a resolution of 512 × 512 pixels. Data augmentation was applied dynamically using the Albumentations library, incorporating geometric transformations (horizontal/vertical flips and transpositions, *p* = 0.5) and photometric perturbations (CLAHE, brightness/contrast adjustments, gamma transformations, hue-saturation shifts, and Gaussian/motion/median blurring, *p* = 0.5), followed by ImageNet normalization. Random cropping was explicitly omitted to maintain consistent spatial reference frames across all samples.

Sub-pixel ground-truth pillar centers were derived from binary masks via an automated geometric estimation routine. Boundary contours were extracted using OpenCV (cv2.findContours), and those meeting minimum structural criteria (area ≥ 80 px^2^, contour points ≥ 40) were fitted with an ellipse via cv2.fitEllipse to yield center coordinates (*c_x_*, *c_y_*), with spatial centroids serving as a fallback for smaller regions. These coordinates were then converted into normalized horizontal and vertical offsets (Δ*x*, Δ*y*) relative to the frame center (*c_xo_*, *c_yo_*) using the formulations 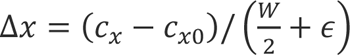 and 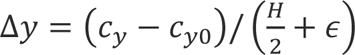 where *W* and *H* represent image dimensions and *ε* = 10^−8^.

The multi-task framework was implemented using a dual-head U-Net architecture with an ImageNet-pretrained ResNet-18 encoder. The segmentation pathway leveraged the standard decoder path to generate single-channel 512 × 512 pixel-wise logits. Concurrently, an auxiliary center regression head was attached to the encoder’s deepest bottleneck features. This regression head applied global adaptive average pooling (AdaptiveAvgPool2d), followed by a 256-unit fully connected layer with ReLU activation and a final linear layer predicting the unconstrained normalized offsets (Δ*x*, Δ*y*).

Model parameters were optimized using a multi-task loss function combining segmentation loss ℒ_seg_ (*W*_seg_ = 1.0) and center regression loss ℒ_center_ (*W*_center_ = 100.0), with the regression loss conditionally applied only to samples containing foreground structures (I_fg_ = 1). The segmentation loss comprised equal weighting of Soft Dice Loss implemented using the MONAI DiceLoss function with sigmoid activation and squared predictions, and Binary Cross-Entropy with Logits, complemented by a small Total Variation regularizer (*w*_TV_ = 2 × 10^−4^) on predicted probabilities to suppress spatial noise. Center regression loss was calculated using Smooth L1 Loss against the normalized ground-truth offsets. Network optimization was conducted over 150 epochs using the Adam optimizer with a learning rate of 1 × 10^−4^, weight decay of 1 × 10^−5^, and a batch size of 20, with the epoch resulting in the lowest total validation loss chosen as the best model checkpoint. Pillar centers during video analysis were determined geometrically from the refined segmentation masks.

### Training of user-specific segmentation models

User-specific models were generated by fine-tuning the pretrained segmentation model rather than training a new model from random initialization. The pretrained U-Net parameters were optimized using the newly annotated images, while the pillar-centre regression component remained frozen. Comparable image normalization and geometric augmentation to the initial model-development pipeline were applied with customizable training duration and batch size.

### Inference pipeline for pillar segmentation (contraction metrics)

Model inference and video evaluation were conducted frame-by-frame across experimental video recordings using identical validation transforms, including reflection padding and resizing to 512 × 512 pixels followed by ImageNet channel normalization. For each extracted frame, predicted single-channel segmentation logits were converted into spatial probability maps via a sigmoid activation function. To suppress high-frequency spatial noise prior to binarization, probability maps were smoothed using a Gaussian filter (7 × 7 kernel, *σ* = 1.2) and thresholded at a probability cutoff of 0.5. The resulting binary masks were then resized to the original video-frame dimensions using nearest-neighbor interpolation prior to morphological post-processing.

To isolate the target pillar structure and remove background artifacts, binary masks were subjected to multi-stage morphological post-processing. Non-contiguous noise was eliminated by retaining only the largest connected component (cv2.connectedComponentsWithStats), followed by sequential morphological closing and opening operations using a 5 × 5 elliptical structuring element. The cleaned binary mask then underwent an iterative refinement routine (*N* = 8 iterations), wherein an ellipse was fitted to the foreground contour (cv2.fitEllipse) and logically intersected with the mask. This process progressively constrained the segmentation boundary, yielding refined sub-pixel center estimates (*c_x_*, *c_y_*) and full major and minor axis lengths (*a*, *b*).

To eliminate frame-to-frame trajectory jitter, estimated center coordinates were smoothed across time using a constant-velocity 2D Kalman filter modeled with a four-dimensional state vector [*x*, *y*, *v_x_v_y_*]*^T^*. Process noise covariances were configured to *Q*_pos_= 2.0 and *Q*_vel_ = 36.0, with a measurement covariance of *R* = 8.0. An outlier rejection gate was implemented to prevent transient tracking failures: raw center predictions exceeding a Euclidean distance of 120 pixels from the filter’s predicted state were rejected, causing the filter to rely strictly on its kinematic motion model for that frame update.

Quantitative kinematic metrics and visual overlays were generated through a two-pass framework. In the first pass, the average ellipse diameter across all valid frames in a video was computed as 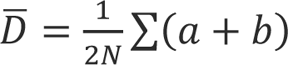 to establish a video-wide, temporally constant bounding circle. In the second pass, diagnostic videos were rendered by drawing this constant bounding circle around the Kalman-smoothed center position onto original video frames, eliminating visual flickering caused by frame-by-frame axis fluctuations. Finally, frame-indexed metrics— including active pillar pixel area, raw and smoothed coordinates, geometric ellipse parameters, Euclidean displacement relative to the video-specific coordinate-wise minimum position (XY_combo_), and area-normalized displacement (XY_combo_norm_) were computed and exported to CSV files for downstream biomechanical analysis (see deep-learning generated overlays in **Video S5**).

### Deep Learning training pipeline for construct segmentation (compaction metrics)

Ground-truth binary masks were generated from manually annotated LabelMe polygons assigned to the *muscle* class. The annotated dataset was randomly partitioned into 85% training and 15% validation subsets using a fixed random seed. To accommodate images acquired at different native resolutions without geometrically distorting the tissues, images and corresponding masks were resized while preserving their original aspect ratio using Albumentations LongestMaxSize, followed by zero-padding to a standardized resolution of 512 × 512 pixels. Training augmentation incorporated horizontal/vertical flipping and transposition (*p* = 0.6), translation (±5%), scaling (±10%) and rotation (±15°; *p* = 0.75), together with photometric perturbations comprising brightness/contrast adjustment, CLAHE or gamma transformation (*p* = 0.9), hue-saturation-value shifts (*p* = 0.3), and Gaussian, motion or median blurring (*p* = 0.25). Images were subsequently normalized using ImageNet channel statistics. Muscle segmentation was performed using a U-Net architecture with an ImageNet-pretrained ResNet-18 encoder and a single-channel segmentation output. The network was optimized using a composite loss comprising MONAI Dice–cross-entropy loss (DiceCELoss; sigmoid activation and squared predictions) and a boundary-sensitive loss weighted by 0.2. For the boundary component, Sobel operators were applied to standardized predicted probability maps and ground-truth masks in the horizontal and vertical directions, and the L1 difference between their resulting gradient magnitudes was minimized to promote accurate delineation of tissue boundaries. Training was performed for 150 epochs using the Adam optimizer with a learning rate of 1 × 10⁻⁴, weight decay of 1 × 10⁻⁵ and a batch size of 32. Validation loss was evaluated after each epoch using the same composite loss, without augmentation, and the network checkpoint yielding the lowest total validation loss was retained for subsequent analysis.

### Deep Learning inference pipeline for construct segmentation (compaction metrics)

For inference, snapshot images underwent the same aspect-ratio-preserving resize, zero-padding and ImageNet normalization used during validation. Segmentation probability maps were generated using sigmoid activation, with horizontal-flip test-time augmentation applied by averaging predictions from the original and horizontally mirrored images after restoring the mirrored prediction to its original orientation. Probability maps were thresholded at 0.50 to obtain binary masks. The masks were refined using sequential morphological closing and opening operations with a 3 × 3 kernel, followed by connected-component filtering. Components with an area of at least 783 pixels at the 512 × 512 network resolution were retained; if no component satisfied this criterion, the largest detected component was retained. Padding was subsequently removed and the refined mask was mapped back to the native image dimensions using nearest-neighbour interpolation, ensuring that morphological measurements were expressed in the original image coordinate system. Muscle compaction-related metrics were calculated directly from the restored binary masks. Tissue area was defined as the total number of foreground pixels. Tissue width was determined independently for each image row containing segmented tissue as the distance between the leftmost and rightmost foreground pixels, inclusive. The mean width across all occupied rows and the minimum row-wise width were calculated for each image. Tissue area, mean width and minimum width, together with the corresponding segmentation metadata, were exported for downstream quantitative analysis.

### Optical flow point tracking algorithm

Point trajectories were estimated between consecutive grayscale video frames using pyramidal Lucas–Kanade optical flow^31^ (cv2.calcOpticalFlowPyrLK; 21 · 21 pixel window, maximum pyramid level of 3, 30 maximum iterations, convergence threshold of 0.01; example tracking **Video S4**; StimTrace workflow: **Video S3**). Initial tracking points were designated either manually or automatically via Shi-Tomasi corner detection. Tracking reliability was verified frame-to-frame using forward–backward consistency,^40^ and trajectory points were accepted only when the round-trip error was ≤ 1.5 pixels. Rejected points were re-evaluated using an expanded recovery search (41 · 41 pixel window, maximum level of 4, 50 maximum iterations). Points failing validation for three consecutive frames were flagged as lost and recorded as missing values (NaN).

### Validation of automated pillar tracking

StimTrace was validated using five recordings (1,136 manually annotated frames; see example of manual segmentation **Video S7**). For every frame, the pillar was independently labelled with an ellipse to establish ground-truth center coordinates and dimensions. The same recordings were analyzed using the raw, unsmoothed ellipse predictions of the deep-learning segmentation model and a manually initialized Lucas–Kanade optical-flow tracker. All recorded displacements were converted from pixels to forces by using identical calibration. Method-specific traces and separate overlay videos were exported for downstream statistical analysis and visual quality control (see StimTrace deep-learning based segmentation **Videos S5,** optical flow point tracking **S4,** manual ground truth segmentation **S7,** and **Figure S6** comparing the different approaches).

### Fabrication of electrostimulation lids

Electrostimulation lids were fabricated by 3D-printing a custom 12-well plate attachment that also served as drill guide to bore holes through the tissue culture plastic lids. Spectroscopically pure graphite rods (SPI, Spectroscopic Pure Graphite Rods for Evaporation) served as electrodes and were cut to a length of 22 mm using a saw guide and additionally a notch was cut into each rod to accommodate the electrical wiring. Silver-plated copper wires (0.6 mm diameter) were attached to the graphite rods using conductive silver epoxy to establish electrical continuity. Temporary aligners maintained uniform electrode spacing while the epoxy cured. The electrodes were potted and sealed into the lid apertures using a biocompatible UV-curable resin (Liqcreate Bio-Med Clear) and polymerized in a UV chamber (BSL-01, Opsytec Dr. Gröbel GmbH). Finally, resistance between electrodes was measured with a multimeter (typically between 2-8 Ω) and a 4 pin connector was soldered on to facilitate easy connection to the stimulator.

### Assembly of electrostimulation device

The lower compartment of the electrostimulation unit housed two Arduino Uno microcontrollers, which controlled the XY-gantry system and rotary stage via dedicated stepper motor drivers. The microcontrollers also operated three dual H-bridge drivers (L298N) to generate six independent, customizable biphasic stimulation channels, as well as status indicator LEDs. To enable serial communication with the host PC software, both Arduinos, the incubator camera, and the digital microscope were interfaced via standard USB connections. The upper unit contained a 220 V AC to 12 V DC power supply; buck converters were integrated with adjustable multi-position switches to generate stimulation potentials ranging from 9 to 15 V DC.

### Fabrication of two-pillar construct

Pillar constructs were prepared by vacuum molding PDMS (base to curing agent ratio of 1:15) into custom-made 3D-printed molds (Prusa SL1s, liqcreate Bio-Med clear resin). To form the rigid pillar, a stiff plastic column was inserted into the PDMS prior to curing and the assembly was cured in an oven at 55°C overnight. After demolding, pillar constructs were rinsed in isopropyl alcohol, air-dried and stored until used. Before the micropillars were used for experiments, cut-outs for the electrodes were punched out the PDMS discs using a 4 mm biopsy punch. Subsequently, the pillar inserts were disinfected with 70% EtOH and dired under UV exposure.

### Cell culture and biofabrication of engineered cardiac tissues

Human peripheral blood mononuclear cells (PBMCs) from a healthy female donor were reprogrammed into induced pluripotent stem cells (iPSCs) using the CytoTune™-iPS Sendai Reprogramming Kit (Thermo Fisher Scientific). iPSCs were maintained ins StemMACS iPSC-BrewMedium (Miltenyi Biotec).^41^ Upon reaching 80% confluence, differentiation into cardiomyocytes was initiated with StemMACSCardioDiffKit XF (Miltenyi Biotec) according to the manufacturer’s instructions. After 10 days, cells were dissociated and used for biofabrication.

Human ventricular cardiac fibroblasts (NHCF-V; Lonza, Cat. CC-2904) were cultured on tissue culture vessels coated with collagen type I (Merck). Cells were maintained in Advanced DMEM (Gibco) supplemented with components of FibroLifeSerum-Free Fibroblast LifeFactors Kit (Lifeline Cell Technology), comprising HSA/linoleic acid/lecithin, FGF, EGF/TGF-β1, insulin, ascorbic acid, L-glutamine, and hydrocortisone hemisuccinate. In addition, fresh FGF was added at each medium change. Cells were cultured until approximately 80-90% confluence and subsequently detached for further processing.

For suspended tissue fabrication, 12-well plates were prepared featuring custom-made PDMS molds. Molds and pillar constructs were sterilized by incubation in 0.5 % peracetic acid in deionized water for 20 minutes at room temperature, followed by extensive rinsing in PBS. Subsequently, molds and pillars were air-dried under UV-light for 60 minutes before the molds were coated with 2 % pluronic F-127 (Sigma-Aldrich) for 1 hour.

For biofabrication, cells (iPSC-CM and cardiac fibroblasts; 75:25 ratio, 10^7^ cells per mL) were mixed with appropriate bovine fibrinogen concentrations (Fibrinogen: 3 - 5 mg/mL in RPMI1640; Sigma-Aldrich, F8630, Thrombin: 0.2 U; Sigma-Aldrich, T4648-1KU). Next, 100 µL of this solution was transferred into the molds, micropillars were put in place and tissues allowed to crosslink in the incubator for 30 minutes without medium. The resulting tissues were cultured in RPMI1640 supplemented with 6-aminocaproic acid (1.5 mg/mL; ACA) fetal bovine serum (5% v/v; Gibco, A5256701), ROCK inhibitor Thiazovivin (2 µM; Stemcell technologies), B27 (2% v/v; Gibco) and primocin (100 µg/mL; InvivoGen). Once completely detached from the mold (usually within 12-48 hours), the cardiac tissues were removed from the molds and transferred on elevation platforms.

Cardiac spheroids were prepared by seeding 4000 cells (iPSC-CM and cardiac fibroblasts; 75:25 ratio) per well of an Akura 96 spheroid microplate. Cells were centrifuged at 250 rcf for 3 min, placed in the incubator at an inclination angle of 30° and allowed to self-assemble.

After the initial 24 hours, the medium was replaced every other day with RPMI1640 supplemented B27 (2% v/v; Gibco) and primocin (100 µg/mL; InvivoGen). All cell cultures and tissue constructs were maintained at 37 °C in a humidified atmosphere containing 5% CO₂ unless otherwise specified.

### Cell culture and biofabrication of engineered skeletal muscle tissues

This study utilized primary human myoblasts obtained from a healthy donor. The line was generated and provided by Dr. Thomas Laumonier (University of Geneva, Switzerland). For this, human muscle biopsies were obtained under approval of the Commission Cantonale d’Éthique de la Recherche of the Canton of Geneva, Switzerland (approved project: “Myogenic stem cells and improvement of muscle regeneration”; approval no. PB_2016-01793; approved on 18 October 2018). Informed written consent was obtained from all study participants in accordance with the guidelines and regulations of the Swiss Health Authorities. All applicable ethical and legal guidelines governing the procurement and use of this cell line were followed. All cell lines were tested for mycoplasma using the MycoStrip™ kit (InvivoGen, rep-mys-10). Human myoblasts were cultured in growth medium in a humidified incubator at 37 °C with 5% CO₂. Myoblast growth medium consisted of DMEM/F-12 supplemented with HEPES (Thermo Fisher Scientific, 11330032), 20% fetal bovine serum (FBS; Thermo Fisher Scientific, 10270106), 1% penicillin/streptomycin (Thermo Fisher Scientific, 15140122), 10 ng/mL human epidermal growth factor (hEGF; Sigma-Aldrich, E9644), 1 ng/mL human basic fibroblast growth factor (human bFGF; Bio-Techne, 233-FB-500), 10 µg/mL human insulin (Sigma-Aldrich, I9278), and 0.4 µg/mL dexamethasone (Sigma-Aldrich, D4902), as previously described.^42^

To induce differentiation, the culture medium was replaced with a low-serum differentiation medium comprising DMEM/F-12 with HEPES, 2% FBS, 1% penicillin/streptomycin, 10 µg/mL human insulin, and 100 µg/mL Normocin (InvivoGen).

For biofabrication, human myoblasts (8·10^6^ cells per mL) were resuspended in a bovine fibrinogen solution (3 - 5 mg/mL in DMEM; 0.2 U Thrombin; 30% v/v Matrigel, Corning, 80094-330;) to generate engineered muscle constructs. For tissue fabrication, 104 µL of the mixture was cast into each mold and micropillars were immediately placed into the precursor and crosslinked in the incubator for 30 minutes without medium. Constructs were cultured in myoblast growth medium supplemented with 6-aminocaproic acid (1.5 mg/mL; ACA) and Normocin (1:500; InvivoGen) for 2 days, after which the culture was switched to low-serum differentiation medium for 3 days prior to electrostimulation.

### Immunofluorescence and confocal imaging

Engineered tissue constructs were incubated with 10 µM blebbistatin for 1 h before fixation to suppress actomyosin contraction and preserve the tissues in a relaxed state for subsequent sarcomere-length measurements. Constructs were then washed with PBS and fixed in 4% (w/v) paraformaldehyde overnight at 4 °C. For permeabilization, constructs were washed with PBS twice and incubated in permeabilization buffer (0.5% v/v Triton X-100 and 5% DMSO in PBS) with gently rocking overnight. Subsequently, tissues were blocked in blocking buffer (1 % w/v bovine serum albumin, 5% v/v normal goat serum and 0.2% v/v Triton X-100 in PBS) for two hours. For primary antibody staining, antibodies were diluted in staining buffer (1 % w/v bovine serum albumin, 1% normal goat serum and 0.05% v/v Triton X-100 in PBS, see **Table 1**) and incubated at 4°C for 24 hours with gently rocking. Samples were washed in washing buffer (0.05% v/v tween-20 in PBS) 3 times for 15 minutes and stained in staining buffer containing secondary antibodies and dyes for 4 hours at room temperature. Before confocal imaging (Fluoview 5000, Olympus), samples were washed extensively in washing buffer.

**Table 1.**
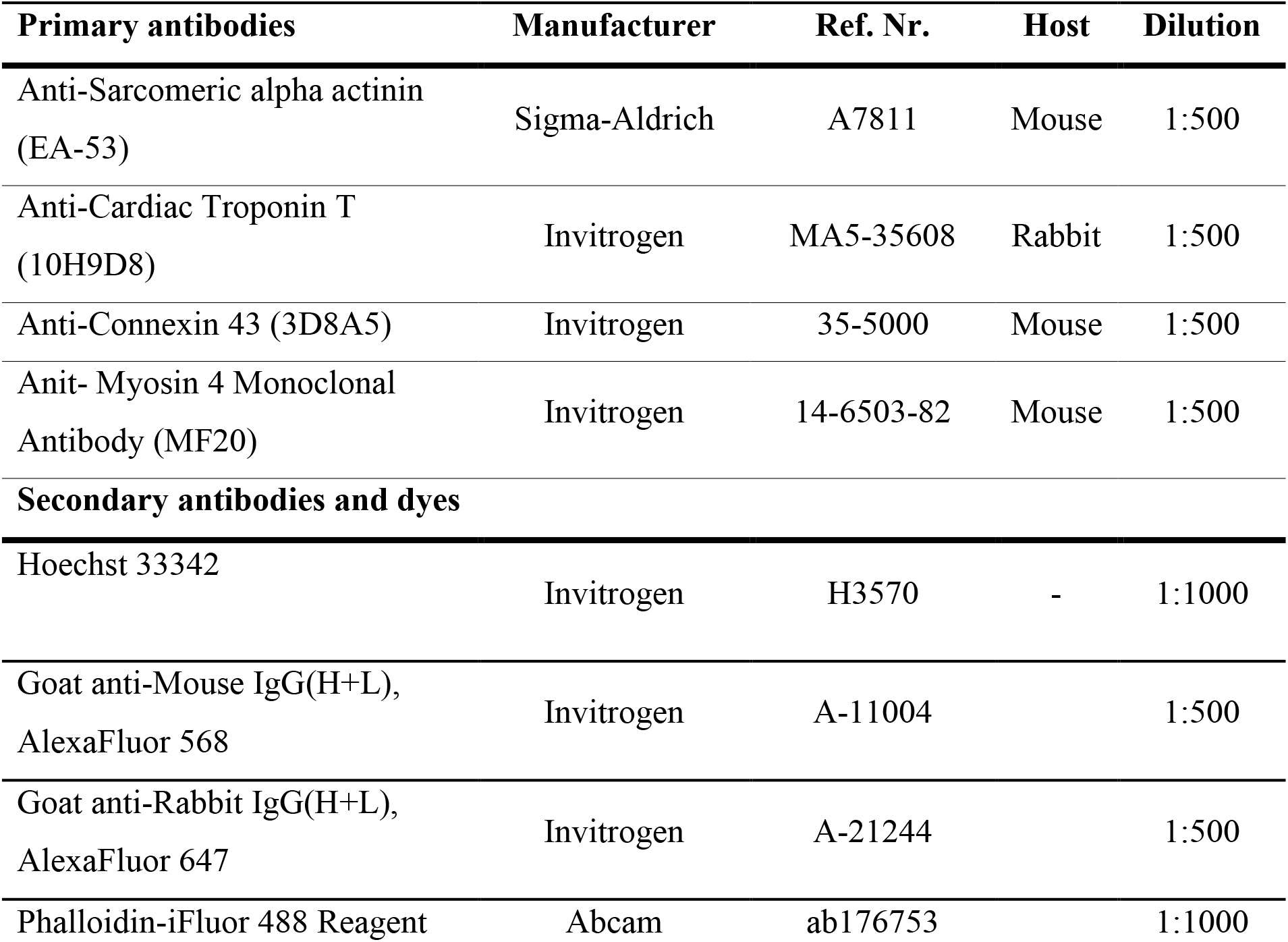
Antibodies and fluorescent dyes dilutions used in study.

### Analysis of immunohistochemical images

Confocal z-stacks were visualized using Imaris (version 10.2, Oxford Instruments), and quantitative morphological analysis was performed using Fiji (ImageJ 1.54p, NIH). For cardiac constructs, sarcomere length was determined from line intensity profiles using the histogram feature, recording 15 measurements per tissue across 3 distinct z-planes. Connexin 43 (Cx43) expression was evaluated by quantifying both total puncta count and individual spot area per region of interest. For skeletal muscle constructs, sarcomere length was evaluated using the same histogram-based line profile protocol. Myotube diameter was determined by measuring the width of myosin heavy chain (MyHC)-positive structures (5 measurements per z-plane across 3 distinct z-levels per tissue). The fusion index was calculated as the ratio of nuclei residing within MyHC-positive multinucleated myotubes to the total number of nuclei. Additionally, F-actin alignment was quantified using the Fiji OrientationJ plugin (available at http://bigwww.epfl.ch/demo/orientationj), applying a structure tensor evaluation with a Gaussian gradient and a local window size of 10 for both distribution and analysis.

### Electrical stimulation

Engineered heart tissues were subjected to electrical stimulation after 7 days of static culture. Graphite electrodes were positioned either parallel to the tissue long axis (collinear configuration) or perpendicular to it (orthogonal configuration). For engineered cardiac tissue conditioning, continuous electrical pulses (8 V biphasic signals corresponding to 2.5 Vcm^-1^, 2 ms per phase, separated by a 1 ms 0 V plateau) were injected. The stimulation frequency was either maintained at 1.5 Hz (continuous) or subsequently increased by 0.5 Hz every other day until reaching 5 Hz (ramping).

Engineered skeletal muscle tissues were subjected to electrical stimulation after 5 days of static culture (2 days in growth medium, 3 days in differentiation medium). Graphite electrodes were placed in collinear configuration. Constructs were exposed to stimulation using one of two regimes: continuous stimulation at 1 Hz (6 min/h) or pulsed stimulation (6 min/h; 1 s trains at 10 Hz followed by a 9 s rest period). Both regimes utilized biphasic square waves of 8 V (2.5 Vcm^-1^); 2 ms per phase separated by a 1 ms 0 V plateau.

### Mechanical testing of PDMS pillars

Pillar mechanics was assessed by fixing the pillar construct to a stable platform before unconfined compression tests were performed using the TA.XTplus Texture Analyzer (Stable Micro Systems) equipped with a 500 g load cell. Samples were placed with the flexible pillar pointing towards the compression plate and a preload of 0.1 g was applied. After relaxation, a displacement of 500 µM was applied at a strain rate of 0.01 mm s−1. Loading and unloading curves were recorded and the force-displacement relationship was established: F(µN) = 6.14·Displacement (µm) + 0.73.

### Finite element method (FEM) of flexible PDMS pillars

Micropillar deflection was simulated in COMSOL Multiphysics (v6.2, COMSOL Inc.) using a 3D solid mechanics finite-element model. The PDMS micropillars and attached tissue were modeled as homogeneous, isotropic, linear elastic materials with Young’s moduli of 700 kPa (measured bulk properties) and 32 kPa, respectively. Fixed constraint boundary conditions were applied to the baseplate and the rigid pillar. To mimic cellular contraction at the tissue attachment interface, a prescribed boundary load was applied to the top region of the flexible pillar. Pillar deflection was evaluated across a force range of 0 to 3000 µN using a stationary parametric sweep.

### FEM simulation of electrical potential

Electric potential and field distributions were simulated in COMSOL using a three-dimensional stationary Electric Currents model reproducing the experimental collinear and orthogonal electrode configurations. The model comprised PBS (electrical conductivity, 1.4 S/m; relative permittivity, 80), PDMS (electrical conductivity, 0.83 × 10⁻¹² S/m; relative permittivity, 2.75), and graphite electrodes (electrical conductivity, 1.28 × 10⁷ S/m; relative permittivity, 12).^36^ A potential difference of 8 V was applied across the graphite electrodes and electric insulation was applied to the micropillars and external faces. The resulting electric potential and electric-field distributions were calculated throughout the device, and the potential along the longitudinal axis of the tissue construct was extracted for comparison with experimental measurements.

### Computation of cardiac metrics

Cardiac metrics were calculated from raw segmentation traces without Kalman filtering or signal smoothing. Beats were identified as local maxima exceeding 20 % prominence and a minimum-interval criteria of 0.1 s was applied. Adjacent local minima defined the pre- and post-beat diastolic values. Time-to-peak was measured from the pre-beat minimum to the maximum of the contraction amplitude and rise 10-90% was the interval between the ascending 10% and 90% amplitude crossings. Maximal contraction and relaxation slopes were the greatest (positive and negative) first derivatives during the corresponding phases. Relaxation and contraction-duration metrics were determined from the descending and ascending amplitude thresholds using linearly interpolated crossing times.

### Statistical analysis

Statistical analysis was performed with GraphPad Prism (v. 10.2.3) using t-tests or one-way ANOVA followed by Tukey post-hoc analysis to compare between groups (a significance level of 0.05 was applied). Significance is indicated as follows: p < 0.05 (*), p < 0.01 (**), p < 0.001 (***), and p < 0.0001 (****); “ns” denotes no significant difference.

## Supporting information

A supporting information document and supplemental videos have been provided by the author.

## Acknowledgements

P.C. acknowledges Ambizione grant PZ00P2_216356 from the Swiss National Science Foundation (SNSF), and support by the Alternatives Research Development Foundation (ARDF Annual Open Grant # 1660730). The authors also acknowledge the ScopeM facility at ETH Zurich for support with microscopy. StimTrace was developed with assistance from OpenAI Codex (OpenAI; accessed August 2026) as an interactive coding tool for implementation, refactoring, and testing. Codex was not used to independently modify experimental data or make scientific decisions. ChatGPT and Google Gemini were used to refine the language. The authors retain full responsibility for the methodology, results, and interpretation.

## Data and source code availability

Source code for StimTrace (the version used in this study) is available at (Stimulatrix: https://github.com/michaelwinkelbauer/Stimulatrix; Stimtrace: https://github.com/michaelwinkelbauer/StimTrace) and permanently archived under DOI https://doi.org/10.3929/ethz-c-000806376. The archived releases include compiled Windows applications, installation instructions, dependency versions, example output and validation materials. Stimulatrix is distributed under the GPL-3.0 license, whereas StimTrace is distributed under the MIT License.

