## Supplementary Figures S1 to S11 for "Stimulatrix: An open-source automated platform for high-throughput functional characterization of engineered contractile tissues"

This file contains **Supplementary Figures S1 to S11**

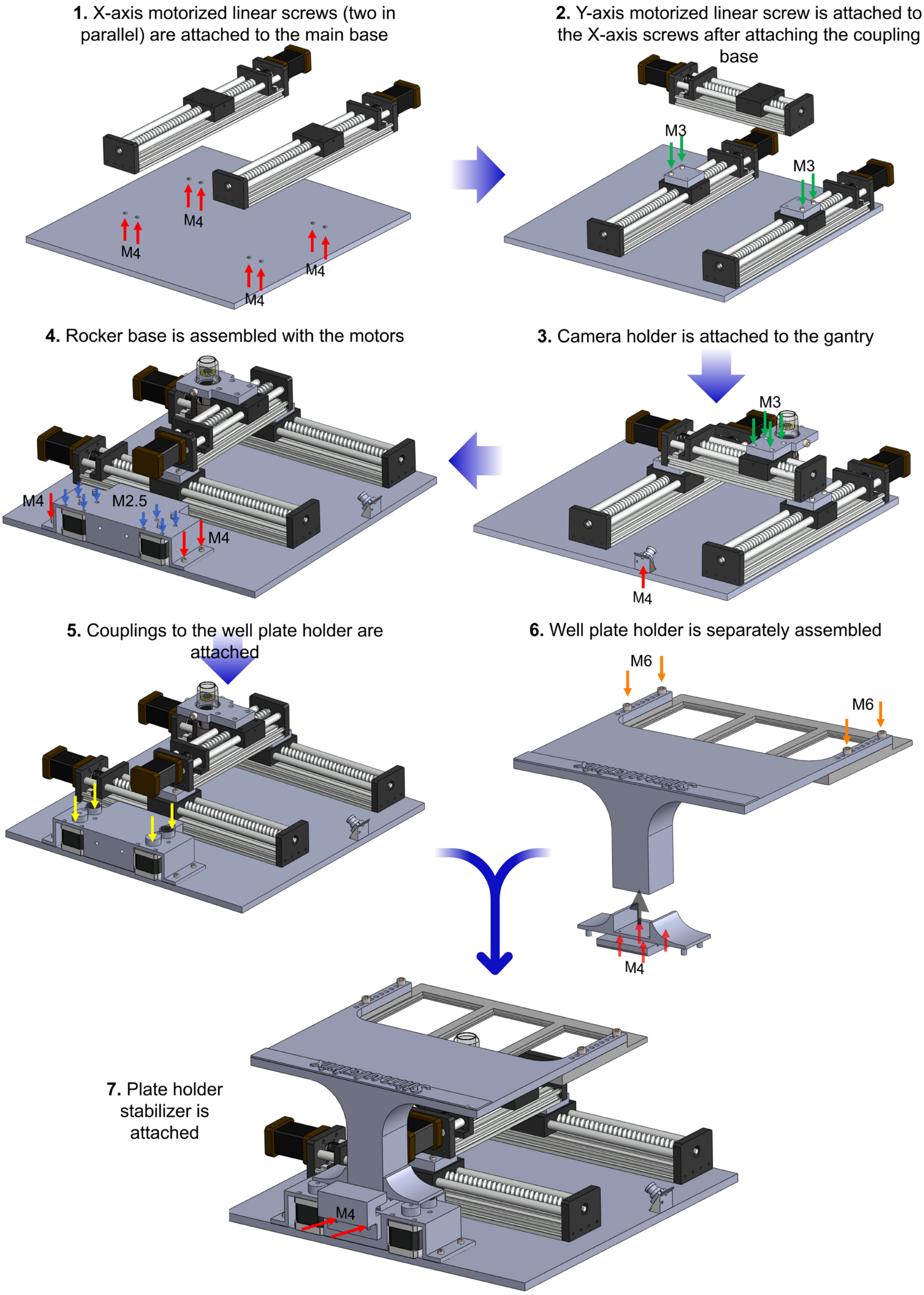


**Figure S1. Assembly steps for the Stimulatrix device.** All components non-commercially-sourced components except the PVC base were 3D printed. **Note:** For each screw positioned mentioned below, a screw insert was added in the absence of counter nut or thread. Here, parts except main base already had the hole positions for putting in the screw inserts.

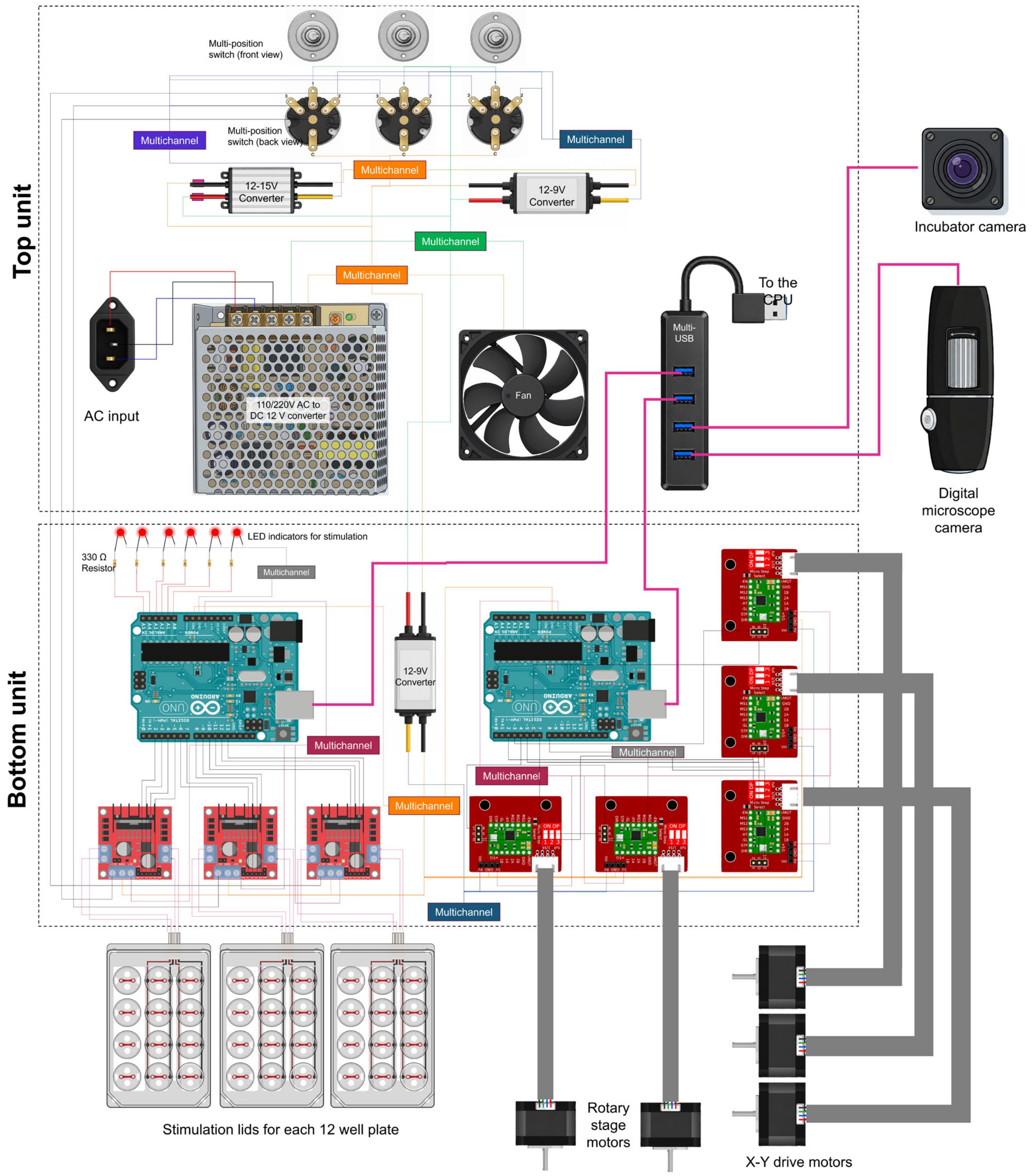


**Figure S2.** Electronic components within the Control Unit of the Stimulatrix Device and their interfacing with the motors, Stimulation lids and Cameras in the Stimulatrix Device.

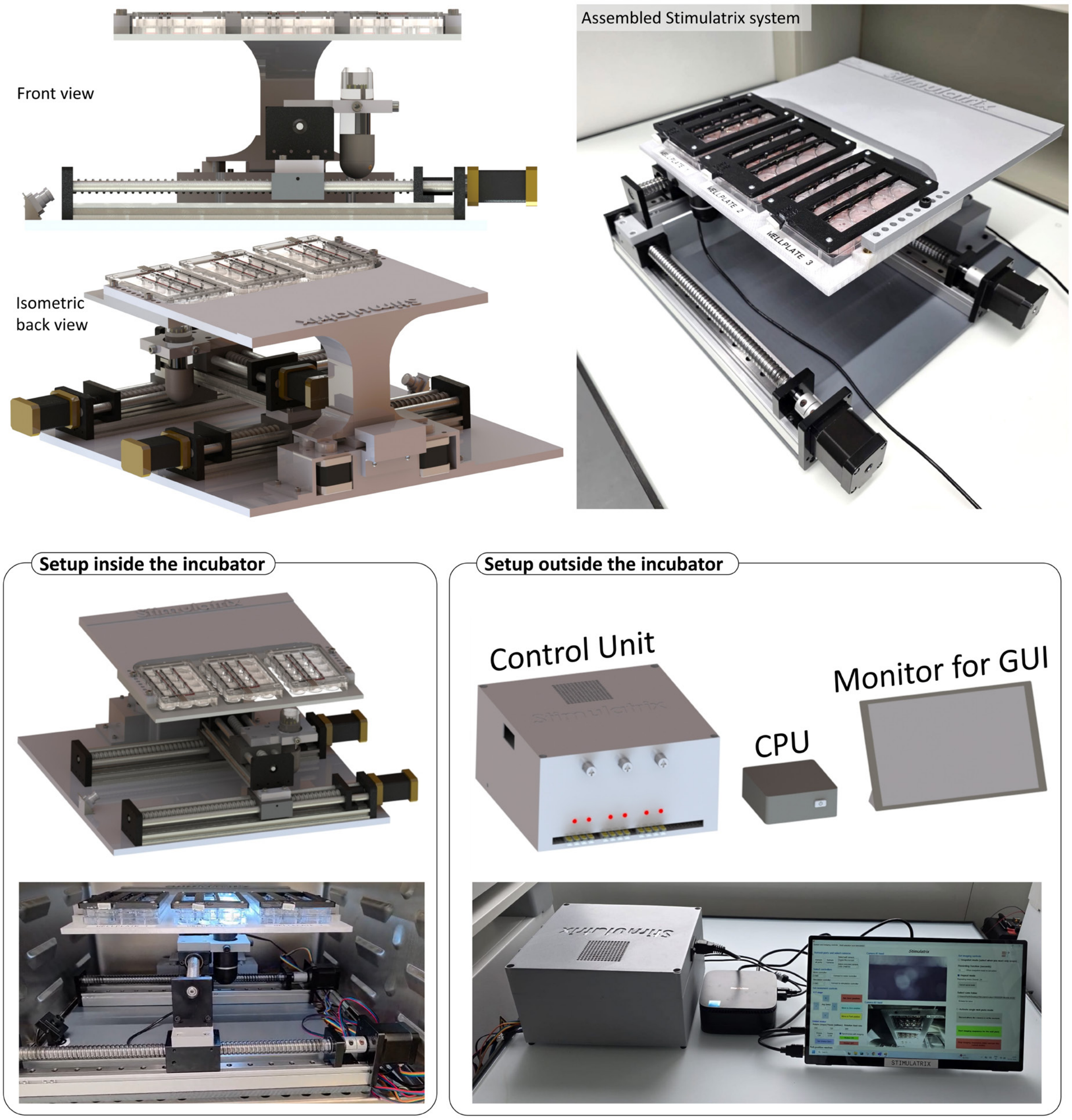


**Figure S3.** Different views of the assembled Stimulatrix system and its associated components.

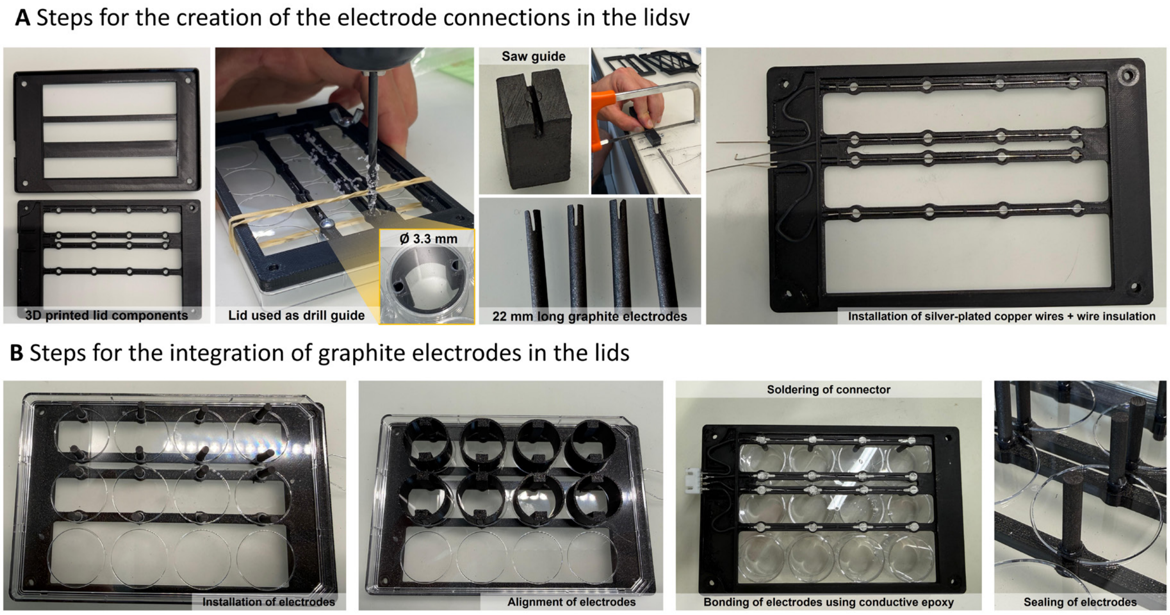


**Fig S4. Fabrication and assembly workflow of the electrical stimulation lid. A.** Custom 3D-printed frame components serve as drill guide to generate aligned holes (Ø 3.3 mm) in standard 12-well plate lids. Graphite rods were cut to length (22 mm) and a slot was cut using a 3D-printed saw guide, which are then integrated with the insulated, silver-plated copper wiring through the 3D-printed frame. **B.** Graphite electrodes are positioned vertically into the slots within the 3D printed lid enclosure and potted with conductive silver epoxy and aligned within the well-plate lid using custom aligners. Electrode feedthroughs and electrical wiring are then fully sealed in biocompatible resin to ensure mechanical stability, fluidic insulation and corrosion resistance during tissue culture.

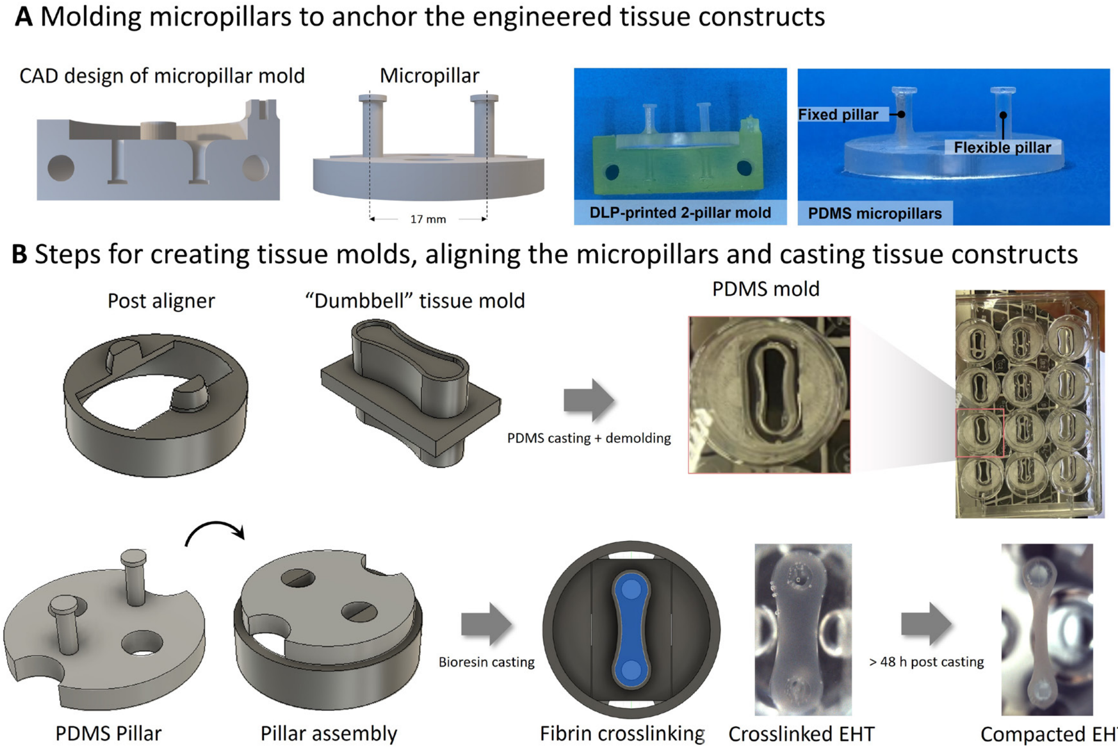


**Figure S5. Mold fabrication, platform assembly, and engineered tissue casting. A.** CAD schematics and photographs of DLP 3D-printed molds used to cast PDMS micropillars with one fixed and one flexible pillar. (Middle) CAD models and representative photos of the dumbel-shaped PDMS tissue molds used in multi-well format. **B.** Step-by-step workflow illustrating pillar assembly with the post aligner, fibrin-based bioresin casting, initial hydrogel crosslinking, and cell-mediated compaction into a 3D engineered tissue (example image shows an engineered heart tissue (EHT)) around the micropillars within 48 h.

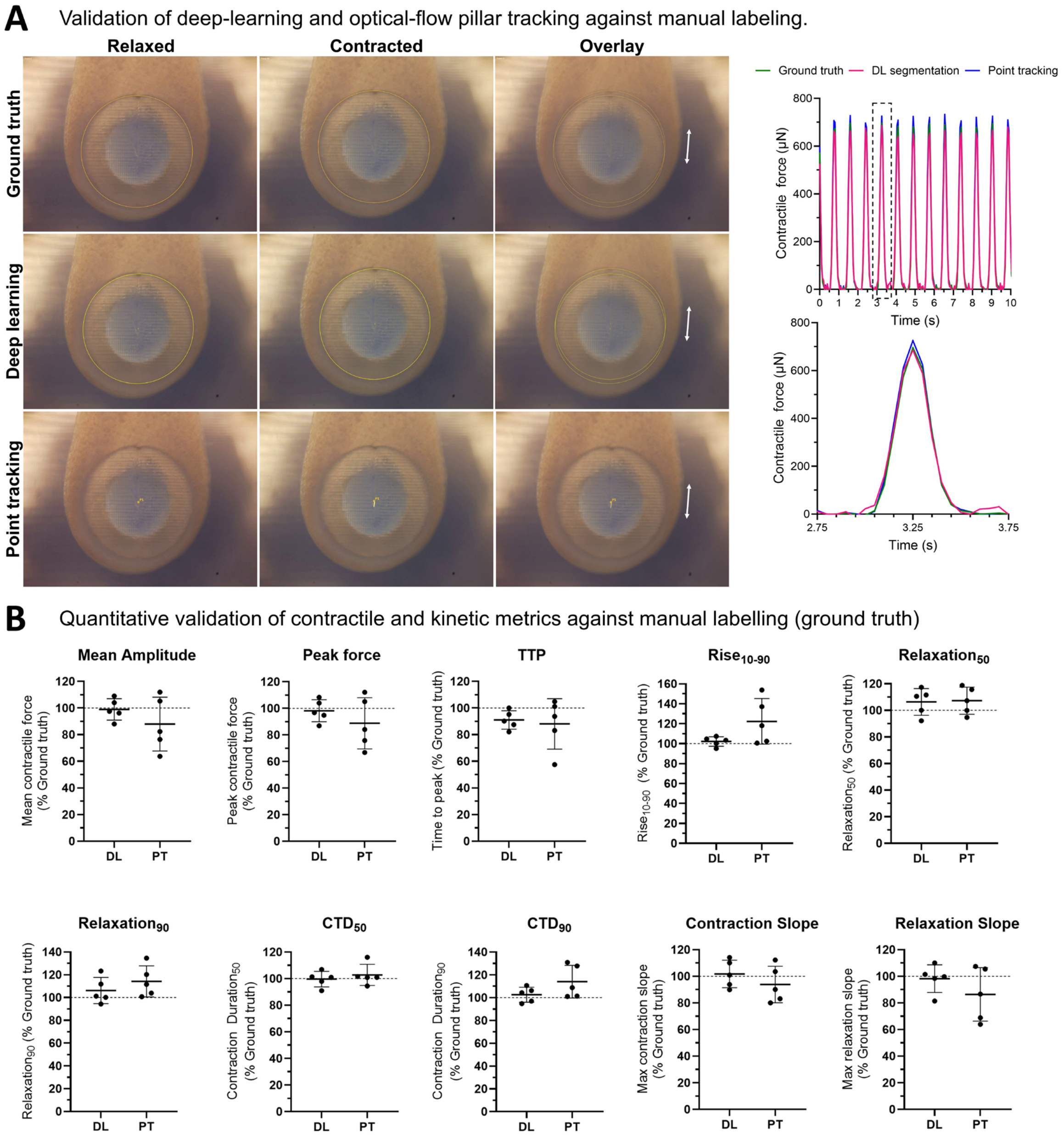


**Figure S6. Validation and performance comparison of automated pillar-tracking algorithms against manual ground truth. A.** Tracking validation and force reconstruction. Representative high-magnification images of micropillars in relaxed, contracted and overlay state analyzed by manual labeling (ground truth; green) deep-learning segmentation (DL; magenta), and optical-flow point tracking (PT, blue). Corresponding continuous force traces over 10 seconds (top right) illustrate high fidelity across all tracking methodologies. **B.** Quantitative comparison of contractile and kinetic metrics normalized to manual labeling. Benchmarking of key metrics extracted via DL or PT, expressed as percentage of manual ground truth (dashed line at 100%). Evaluated parameters include amplitude metrics (mean amplitude, peak force), kinetic durations (time to peak (TTP), $Rise_{10-90}$, $Relaxation_{50}$, $Relaxation_{90}$, contraction $duration_{50}$ ($CTD_{50}$), contraction $duration_{90}$ ($CTD_{90}$) and contraction and relaxation slopes). DL segmentation exhibits higher accuracy and reduced variance across all parameters when compared to tracking.  Data presented as individual data points with mean ± SD; n=5 independent recordings of contracting tissues.

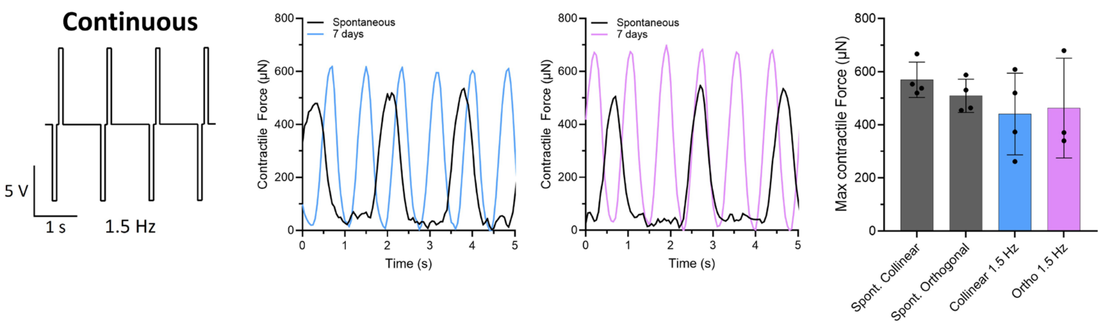


**Figure S7. Long-term electrical pacing regimen in cardiac tissues.** Schematic of the continuous pacing protocol (1.5 Hz over 14 days) with corresponding force traces before and after 7 days of conditioning. Bar chart shows maximum developed contractile force across spontaneous and paced conditions in collinear vs. orthogonal configuration.

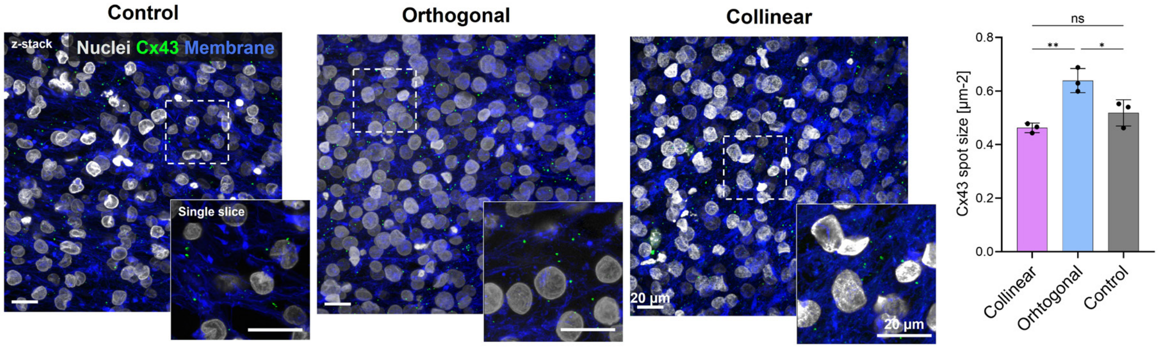


**Figure S8. Characterization and quantification of Connexin 43 (Cx43) spot size.** Maximum intensity z-stack projections and single-slice insets (dashed boxes) showing nuclei (white), cell membrane (blue), and Cx43 (green (scale bars: 20 µm). Bar graph shows quantitative analysis of mean Cx43 spot size across conditions (mean $\pm$ SD, $n = 3$). Statistical significance was determined by one-way ANOVA with Tukey's post-hoc test and is denoted as follows: *p < 0.05, **p < 0.01, ns = not significant).

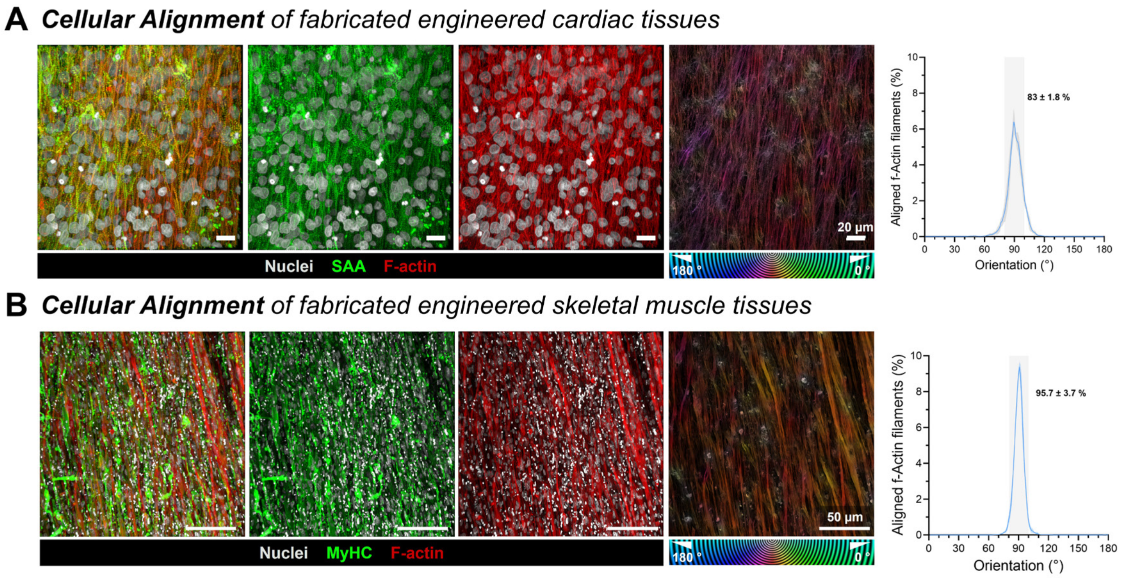


**Figure S9. Analysis of f-actin alignment. A.** The summed proportion of aligned actin filaments within ± 10° of the dominant orientation peaks in engineered cardiac tissues. **B.** The summed proportion of aligned actin filaments within ± 10° of the dominant orientation peaks in engineered skeletal muscle tissues.

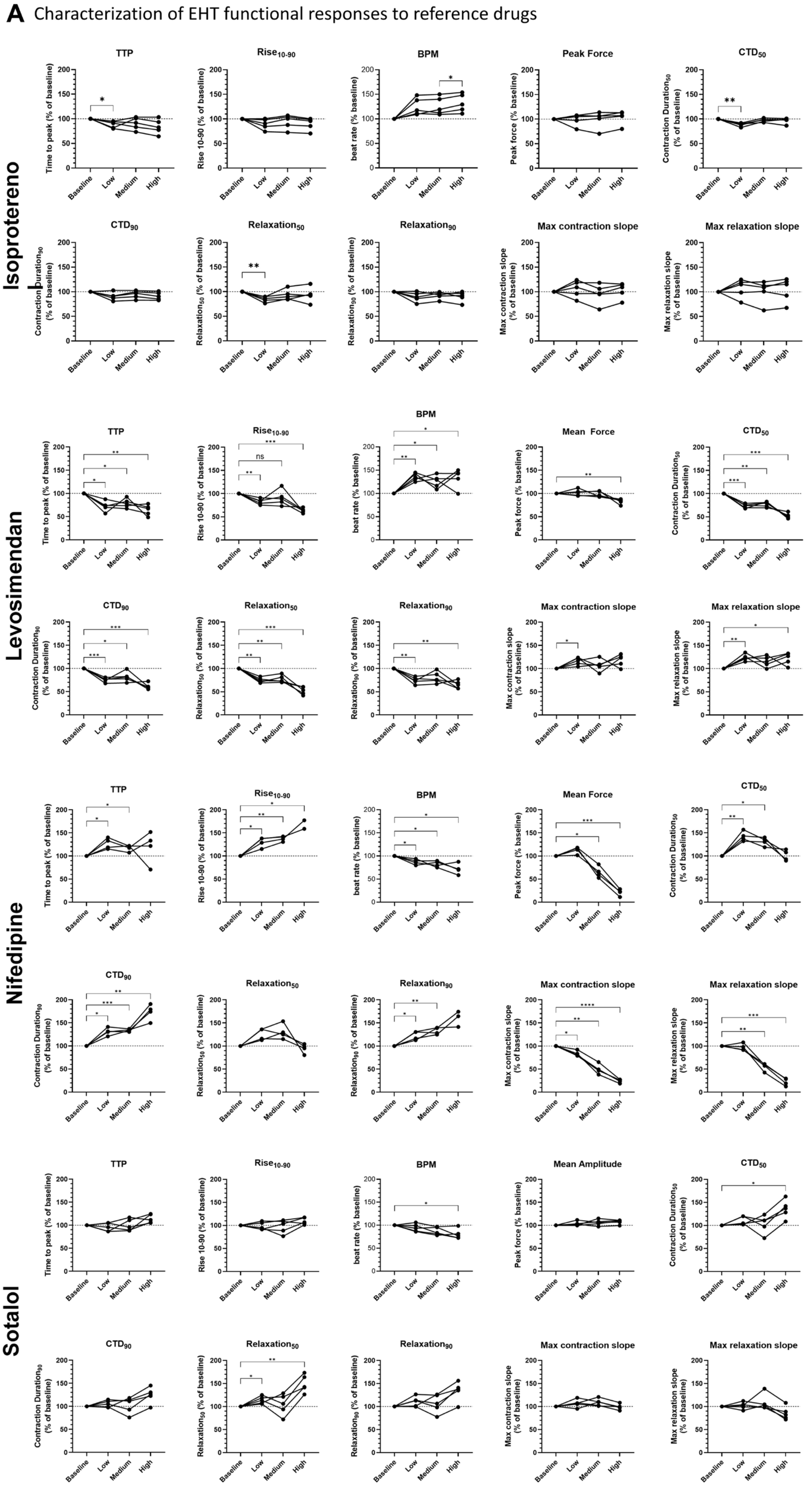


**Figure S10. Dose-dependent functional and kinematic characterizations of cardioactive compounds.** Quantitative analysis of contractile and kinetic metrics in engineered cardiac tissues exposed to increasing concentrations (Baseline, Low, Medium, High) of Isoproterenol, Levosimendan, Nifedipine, and Sotalol. Evaluated parameters include time to peak (TTP), 10–90% rise time ($Rise_{10\text{-}90}$), beat rate (BPM), peak/mean force or amplitude, contraction durations ($CTD_{10}$ and $CTD_{90}$), relaxation durations ($Relaxation_{50}$ and $Relaxation_{90}$) and maximal contraction and relaxation slopes. All metrics are expressed as percentage of individual baseline values (100%, horizontal dashed line). Connected line points represent individual tissue trajectories across concentration steps (n = 5 independent biological replicates). Statistical significance was determined by one-way ANOVA with Tukey's post-hoc test and is denoted as follows: *$p < 0.05$, **$p < 0.01$, ***$p < 0.001$, ****$p < 0.0001$).

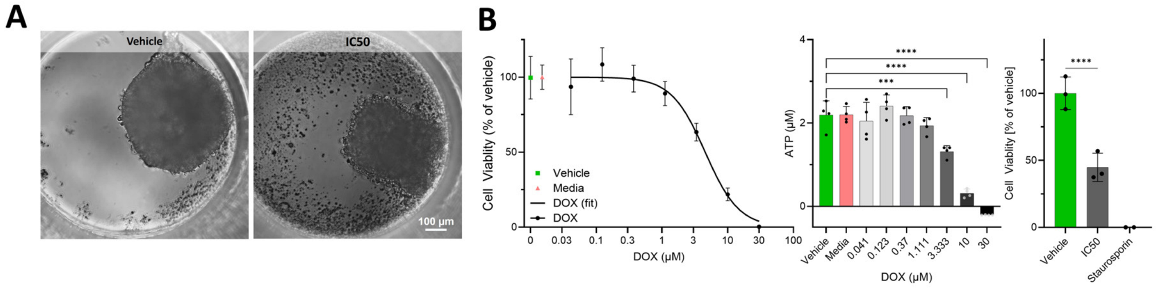


**Figure S11. Assessment of doxorubicin-induced cytotoxicity and $IC_{50}$ determination in cardiac spheroids following 48 h exposure.** (A) Representative brightfield images of cardiac spheroids treated with vehicle control or doxorubicin at the calculated $IC_{50}$ concentration after 48 h (scale bar: 100 µm). (B) Dose-response curve of doxorubicin (0.03 - 30 µM) showing relative cell viability (% of vehicle; left), ATP content (µM; middle), and comparative cell viability relative to staurosporine (1 µM; right) after 48 h incubation. Data presented as mean ± SD (n = 4 independent replicates). Statistical significance was determined by one-way ANOVA with Tukey's post-hoc tests and is denoted as follows: *** $p < 0.001$, **** $p < 0.0001$).
